# Interchromosomal contacts between regulatory regions trigger stable transgenerational epigenetic inheritance in Drosophila

**DOI:** 10.1101/2023.07.13.548806

**Authors:** Maximilian H. Fitz-James, Gonzalo Sabarís, Peter Sarkies, Frédéric Bantignies, Giacomo Cavalli

**Affiliations:** Institute of Human Genetics, CNRS and University of Montpellier, 141 Rue de la Cardonille, 34094 Montpellier, France; Department of Biochemistry, University of Oxford, South Parks Road, Oxford OX1 3QU, UK

**Keywords:** Transgenerational Epigenetic Inheritance, Chromatin Contacts, Genome Organisation, Epimutation, Polycomb, GAGA-Factor, *Fab-7*

## Abstract

Non-genetic information can be inherited across generations by the process of Transgenerational Epigenetic Inheritance (TEI). TEI can be established by various triggering events, including transient genetic perturbations. In *Drosophila*, hemizygosity of the *Fab-7* regulatory element triggers inheritance of the histone mark H3K27me3 at a homologous locus on another chromosome, resulting in heritable epigenetic differences in eye colour. By mutating transcription factor binding sites within the *Fab-7* element, we demonstrate the importance of two proteins in the establishment and maintenance of TEI: GAGA-factor and Pleiohomeotic. We show that these proteins function by recruiting the Polycomb Repressive Complex 2 and by mediating interchromosomal chromatin contacts between *Fab-7* and its homologous locus. Finally, using an *in vivo* synthetic biology system to induce them, we show that chromatin contacts alone can establish TEI, providing a mechanism by which hemizygosity of one locus can establish epigenetic memory at another *in trans* through long-distance chromatin contacts.

## Introduction

Epigenetic information has long been known to be a major factor in the regulation of gene expression^1^. Whether such information can be transmitted across generations in various organisms has been a more elusive question, made difficult by the potential for genetic factors to confound experiments on heredity^2,3^. Through careful experimentation in model organisms, recent work has now demonstrated that such transgenerational epigenetic inheritance (TEI) does occur in a variety of organisms^4,5^. In addition, although much more work is required, the molecular mechanisms underlying these instances of inheritance have begun to be described^6^. Many of the most well-known epigenetic regulators of gene expression have been implicated in different cases of TEI, including non-coding RNAs^7–12^, DNA methylation^13–15^ and histone modifications^12,16–19^, while less typical sources of non-genetic information, such as 3D chromatin organisation^19^ and transcription factor binding^20^, have been suggested as secondary signals in some cases.

Just as genes and their allelic variants are the basis of genetic variation, so are “epialleles” the basic units of heritable epigenetic change. Similarly, while mutation is the means by which genetic variation arises, “epimutation” describes the appearance of a heritable change in epigenetic information that gives rise to an epiallele. Epimutation provides an alternative source of heritable variation which differs from genetic mutation in that it has the potential to be more rapid, targeted and reversible, allowing for fast adaptation to a fluctuating environment^21^. Nonetheless, the underlying causes of epimutation and the mechanism by which they arise remain unclear and likely vary between organisms.

Given the complexity of mechanisms involved in TEI in higher eukaryotes, model systems that can be easily manipulated and allow one to both effectively track heritable phenotypes and analyse the underlying molecular events at play are critically needed in the field. One clear example of TEI occurs in *Drosophila melanogaster*, in a transgenic line called “Fab2L”, involving a transgene that drives the expression of the eye pigmentation gene *mini-white*^22^. Flies of the Fab2L line exhibit a stochastic phenotype, manifesting as mosaicism of pigmentation in the adult eye. A memory of this phenotype can be established by a transient genetic perturbation^19^ which can then be maintained epigenetically for countless generations following the initial trigger. While this represents a clear case of TEI, its molecular mechanisms remain mysterious, making it a valuable model system to study the means by which heritable epigenetic variability arises. Here, we investigate the mechanistic basis for the establishment of transgenerational epigenetic inheritance at the Fab2L locus. We show that it is mediated by two key regulatory regions of the transgene through the binding of the transcription factors GAGA-factor (GAF) and Pleiohomeotic (Pho). We show that these transcription factors recruit the Polycomb Repressive Complex 2 (PRC2) to the transgene, leading to deposition of H3K27me3, and promote interchromosomal chromatin contacts between the transgene and a homologous region elsewhere in the genome. Using an *in vivo* synthetic biology system, we artificially recapitulate these contacts to demonstrate that chromatin contacts alone are sufficient to induce TEI in *Drosophila*, providing a mechanism whereby genetic perturbation of one locus can trigger TEI at another *in trans* through long-distance chromatin interactions.

## Results

### Binding of GAGA-Factor and Pho is responsible for epigenetic variability at the Fab2L transgene

The *Drosophila* Fab2L line carries a single copy 12.4kb transgene inserted into chromosome arm 2L at cytogenetic position 37B^22,23^. This transgene contains the reporter genes *LacZ* and *mini-white* under the control of the *Fab-7* element. *Fab-7* is a well-studied regulatory region of the bithorax complex on chromosome 3, where it regulates the expression of the Hox gene *Abd-b*^24–26^. Importantly, the Fab2L line therefore contains two versions of the *Fab-7* element in its genome, one at its endogenous location on chromosome 3 and one inserted ectopically within the transgene on chromosome 2 (Figure 1A).

**Figure 1.**
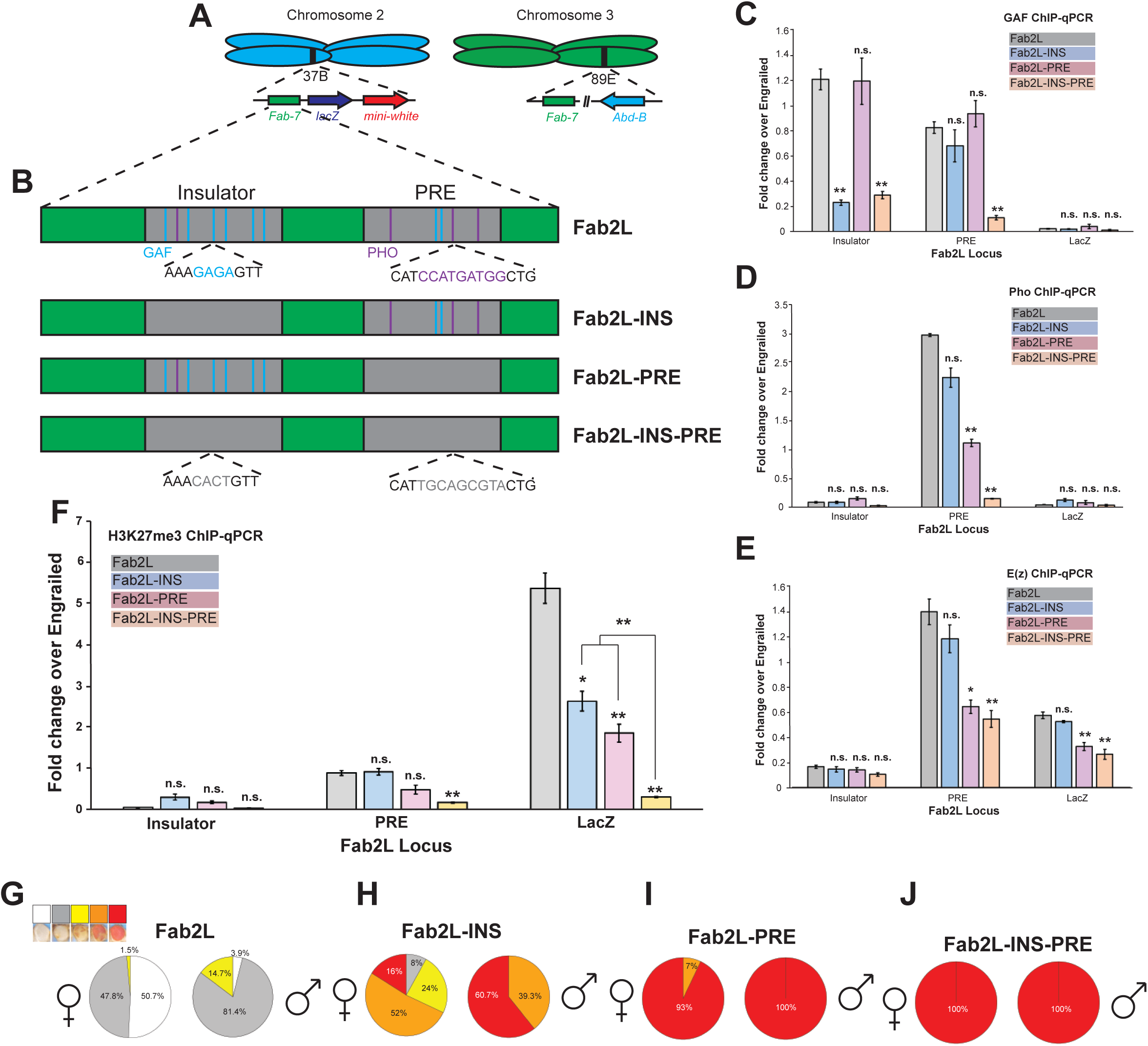
Mutation of GAF and Pho binding sites decreases epigenetic variability of the Fab2L transgene. **(A)** Schematic representation of the Fab2L transgene at its insertion site at cytological position 37B on chromosome 2, alongside the homologous *Fab-7* region on chromosome 3. **(B)** Illustration of the *Fab-7* element with important subdomains and transcription factor binding sites in wild-type and mutated versions of the Fab2L transgene. **(C-F)** ChIP-qPCR assays performed in embryos of the indicated genotypes at regions within the Fab2L transgene. Error bars represent +/- standard error from the mean (SEM) of three independent repeats. Samples were normalised to engrailed as a positive control and compared to wild-type Fab2L or between each other by the t-test (*p<0.05, **p<0.01, n.s. = not significant). **(G-J)** Phenotypic classification of eye colour in female and male adults of the indicated genotypes. Flies were sorted into five classes on the basis of eye colour, representing the number of pigmented ommatidia: Class 1 = 0% ; Class 2 = 1-10% ; Class 3 = 10-75% ; Class 4 = 75-99% ; Class 5 = 100%. See also **Figure S1**.

The *mini-white* reporter gene, which controls red pigment deposition in the eye, is not expressed uniformly in Fab2L flies but shows a mosaic pattern of eye pigmentation, with some ommatidia showing strong *mini-white* expression and others strong repression, within the same individual (Figure S1A,B). Stochastic binding of the Polycomb Repressive Complex 2 (PRC2), resulting in deposition of the repressive histone mark H3K27me3, has been suggested as an explanation for the variability of mini-white expression in transgenes carrying Polycomb-bound elements. This mosaic eye pattern therefore represents a very evident and visible instance of epigenetic variation in the absence of any underlying genetic change^19^.

To further investigate the mechanism behind this epigenetic variation, we generated transgenic versions of the Fab2L transgene with mutations in key sequence motifs in the *Fab-7* element. *Fab-7* contains within it two important subdomains: an insulator region, which in its endogenous state prevents mis-regulation of *Abd-B* by adjacent regulatory regions in the wrong body segments, and a Polycomb Response Element (PRE), which recruits either Polycomb Group (PcG) or Trithorax Group (TrxG) proteins to maintain the pattern of *Abd-B* expression established early in development. Both of these regions contain several consensus sequence motifs for the DNA-binding proteins GAF and Pho (Figure 1B). Among other functions, both of these proteins are known to be recruiters of PRC2, strongly suggesting their potential involvement in the variable H3K27me3 levels, and thus the eye colour phenotype, in Fab2L.

We generated three transgenic lines mutating all GAF and Pho sequence motifs within either the insulator (Fab2L-INS), the PRE (Fab2L-PRE) or both (Fab2L-INS-PRE) in the Fab2L transgene (Figure 1B). Fab2L-INS showed a significant decrease in GAF binding to the insulator, although Pho binding was little affected as was GAF binding to the PRE. Interestingly, Fab2L-PRE had the opposite effect, with GAF binding at both sites unaffected, while Pho binding was significantly decreased at the PRE only (Figure 1C,D). These results extended to the recruitment of PRC2 to Fab2L, which was largely unaffected in Fab2L-INS but significantly decreased in Fab2L-PRE, not only at the PRE itself but also exterior to the *Fab-7* at the downstream LacZ region (Figure 1E). Mutation of both regions together shows clear additive effects. Indeed, in the Fab2L-INS-PRE line binding of both GAF and Pho are decreased at both the insulator and PRE, and in most cases to a significantly greater extent that in either of the single mutants (Figure 1C,D), although PRC2 binding was not significantly different from Fab2L-PRE (Figure 1E). Taken together, these results suggest that the insulator and PRE regions of *Fab-7* act cooperatively to recruit GAF and Pho to the Fab2L transgene, although the PRE may play the greater role in the subsequent recruitment of PRC2.

We then analysed the downstream effects of this altered GAF and Pho recruitment on the chromatin and phenotype of the mutant lines. All three mutant lines had significantly decreased levels of H3K27me3 across the transgene, with the decrease being much more pronounced in the double mutant Fab2L-INS-PRE line (Figure 1F). These changes in chromatin translated to phenotypic effects on the eye colour of the adult flies. Indeed, all three mutant lines displayed shifts towards red eye colour compared to naïve Fab2L (Figures 1G-J and S1). While it remained considerable, the shift in Fab2L-INS was milder than for the other two lines, with approximately 16% of females and 61% of males exhibiting fully red eyes (Figures 1H and S1C,D). In contrast, almost all Fab2L-PRE and Fab2L-INS-PRE flies of both sexes exhibited fully red eyes. It is interesting to note, however, that while the shift towards red was complete for Fab2L-INS-PRE, with all individuals having uniform red eyes (Figures 1J and S1G,H), Fab2L-PRE retained some stochasticity, with around 7% of females possessing at least some white ommatidia (Figures 1I and S1E). These results reinforce the idea that the insulator and PRE together are responsible for the epigenetic and phenotypic variability of the Fab2L fly line.

### The insulator and PRE regions of *Fab-7* are individually sufficient to mediate epigenetic inheritance

In WT flies carrying the Fab2L transgene, epigenetic differences in expression between individuals are not inherited transgenerationally under normal conditions.. Indeed, when we applied repeated selection and crossing of the most extreme individuals in the population of an unmanipulated Fab2L line over ten generations, we did not obtain any significant differences in eye colour across the population (Figure S2A,B). However, crossing this “naïve” Fab2L line with another Fab2L line bearing a homozygous deletion of the endogenous *Fab-7* gives F1 individuals carrying the transgene in a homozygous state, but the endogenous *Fab-7* in a hemizygous state (Figure S2C). This hemizygosity establishes transgenerational epigenetic memory at the transgene, such that reconstituting the Fab2L genotype in the F2 results in a line which is genetically identical to the P0 Fab2L, but in which TEI is now possible^19^. Indeed, selection of this line over 10 generations resulted in either red or white “epilines”: populations of flies with a significant proportion of individuals with monochrome eye colour, i.e. with 100% of their ommatidia either pigmented or unpigmented (Figure S2D). We also found that other crossing schemes which induced *Fab-7* hemizygosity for one or two generations, were able to trigger TEI in Fab2L (Figure S2E-J).

Given the altered phenotype of the mutant Fab2L transgenic lines, we asked whether these mutations also interfered with the ability of the Fab2L transgene to maintain a memory of its epigenetic state across generations by TEI. Due to the shift towards red eyes in the naïve Fab2L-INS, Fab2L-PRE and Fab2L-INS-PRE lines, selection towards red eyes would not be informative. We therefore performed a transgenerational epigenetic selection experiment to determine if these mutant lines could be selected towards a more repressed, white-eyed phenotype than the naïve population. Just as with wild-type Fab2L (Figure S2C), we introduced a single generation of *Fab-7* hemizygosity while leaving the mutant versions of the Fab2L transgene unmanipulated (Figure 2A). We then reconstituted the parental genotype, homozygous for the endogenous *Fab-7*, and selectively bred the most white-eyed individuals over subsequent generations. As a control, we used a wild-type Fab2L line which had previously been selected for a red-eyed phenotype. The Fab2L, Fab2L-INS and Fab2L-PRE, which all showed a greater or lesser degree of variability in their starting populations, were receptive to selection, showing a clear and gradual shift towards whiter eyes in both females and males over the generations (Figures 2B-D and S3A-C). In the case of Fab2L-INS, the appearance of some individuals with fully white eyes was even observed (Figures 2C and S3B), consistent with the less extreme de-repression observed in this line compared to the other mutants. In contrast, the Fab2L-INS-PRE line displayed no variation at any point during the experiment, with all individuals of both sexes maintaining a uniform red eye colour (Figures 2D and S3C). As expected, some pre-existing degree of Polycomb binding and some form of epigenetic variation at the Fab2L transgene is therefore prerequisite for TEI. However, these results also demonstrate that the insulator or PRE regions alone are still able to mediate TEI, indicating that they act together to maintain an epigenetic memory at the transgenic *Fab-7* element in the Fab2L line.

**Figure 2.**
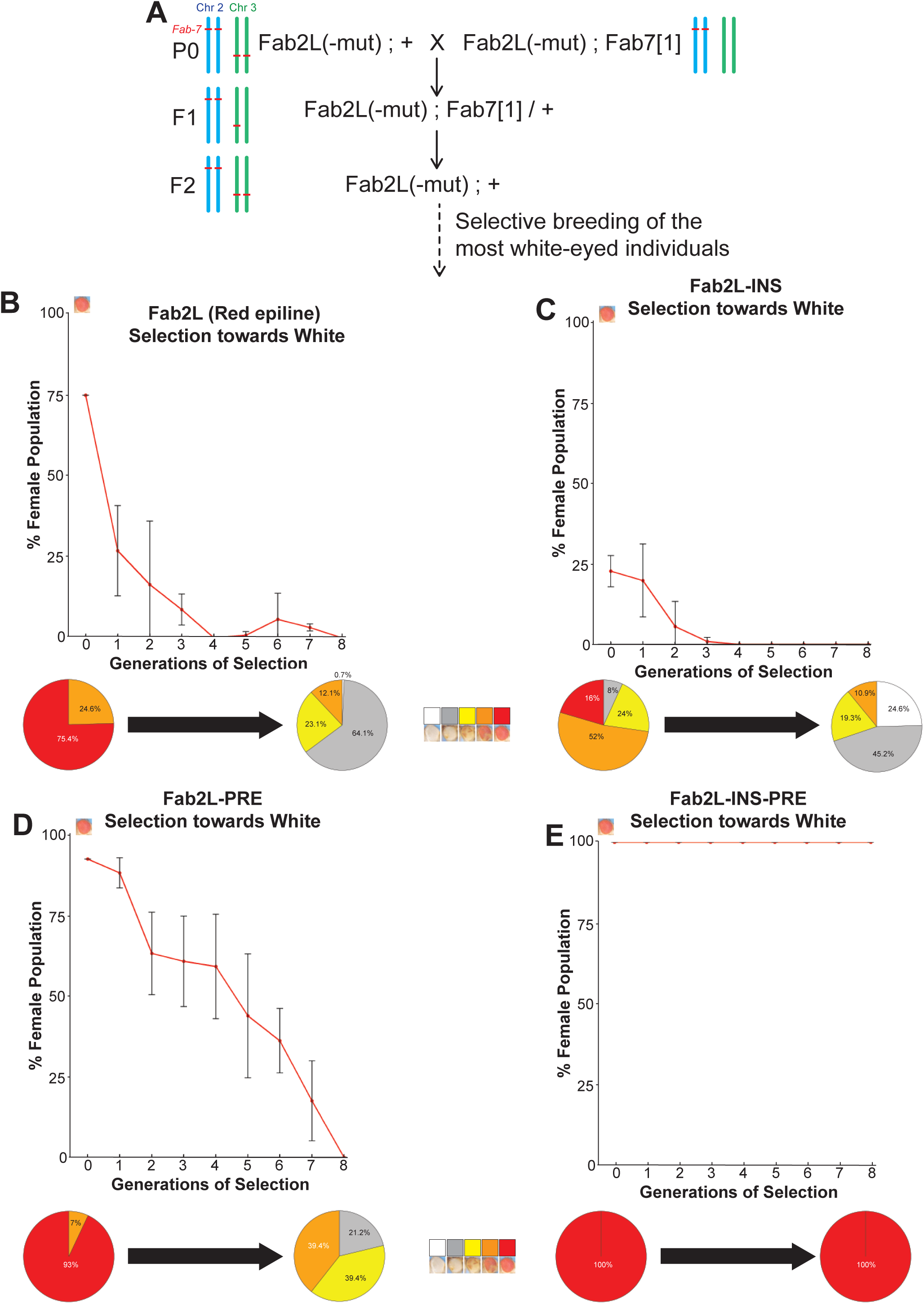
Epigenetic inheritance of eye colour is abrogated in the absence of GAF and Pho binding to the Fab2L transgene. **(A)** Crossing scheme for the triggering of TEI at wild-type and mutant versions of Fab2L, with diagrammatic representation of the copy number of the *Fab-7* element on chromosomes 2 and 3. See also **Figure S2**. **(B-E)** Results of selection for the most white-eyed flies in each generation beyond the F2 of the crossing scheme. At top, curves represent the percentage of Class 5 females in the population across generations. Error bars are +/- standard deviation (SD) of 3 independent repeats. At bottom, pie charts represent the phenotypic distribution of the eye colour within the population in the first and last generations of selection, sorted into five classes. See Also **Figure S3**.

### The insulator and PRE regions of *Fab-7* are required for horizontal transmission of a repressed epigenetic state through paramutation

The Fab2L transgene is not only able to acquire an altered epigenetic state by selection over generations, but can do so in a single generation by the process of “paramutation”^19^. Paramutation denotes the horizontal transfer of an epigenetic state *in trans* between two homologous alleles, and has been described in many organisms including *Drosophila*^27–30^. In the Fab2L line, crossing a naïve Fab2L with an established Fab2L epiline (either white or red-eyed) results in the acquisition by the naïve allele of the altered epigenetic state of the epiline allele. This phenomenon can be tracked by the use of a *black[1]* marker allele, closely linked to the Fab2L transgene, such that F2 individuals that have inherited both copies of Fab2L from the naïve parent can be determined with high probability (Figures 3A and S4A). Although these F2 flies possess the genetic material of the naïve P0 population, their epigenetic state resembles that of the epiline with which it was crossed, attesting to the acquisition over this genomic region of a new epigenetic state (Figures 3B and S4B).

**Figure 3.**
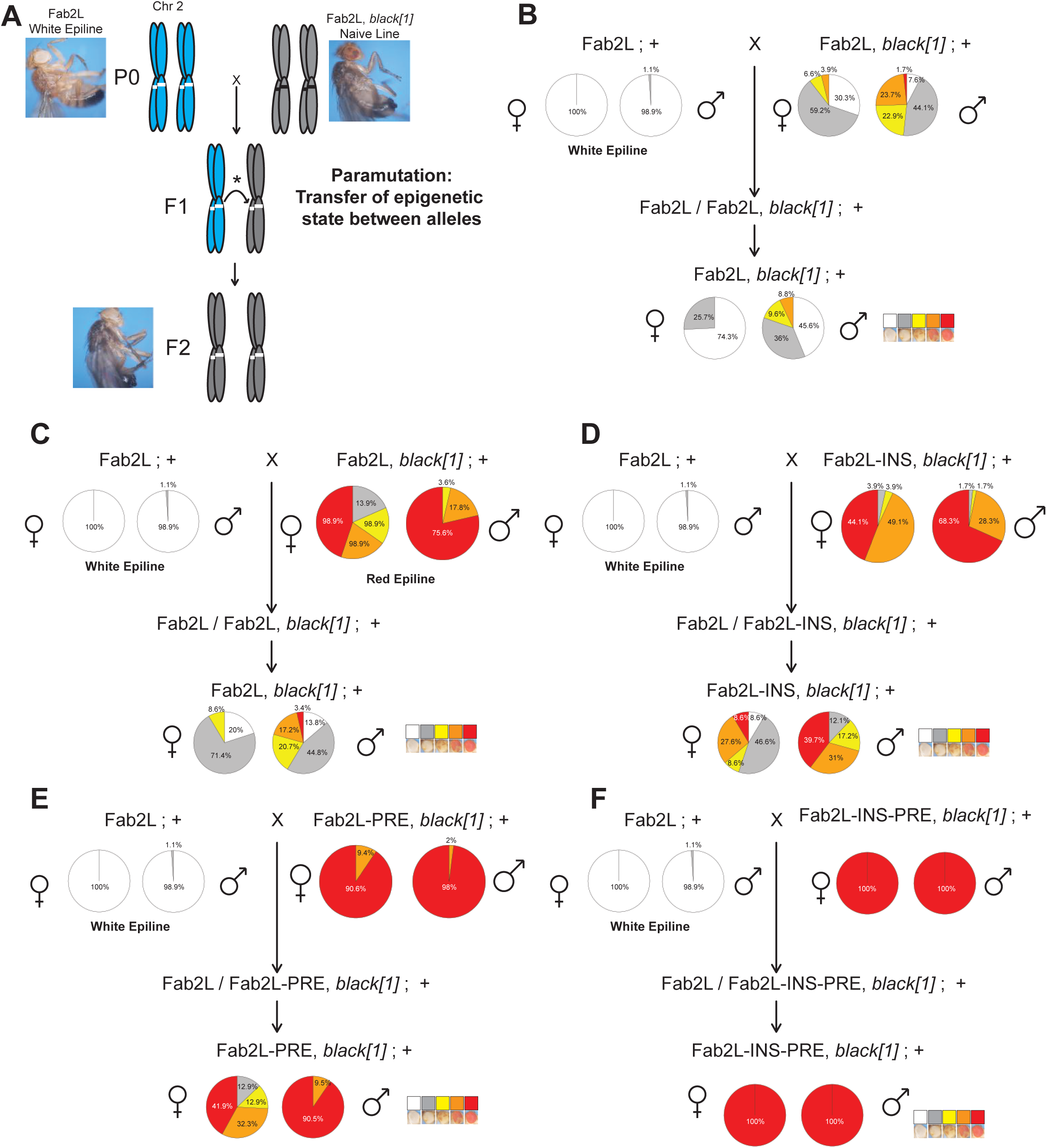
The insulator and PRE regions are required for Fab2L to acquire a repressed epigenetic state through paramutation. **(A)** Illustration of the paramutation crossing scheme for the acquisition of a repressed epigenetic state by a naive Fab2L allele from an established epiallele *in trans*. The presence of a *black[1]* marker linked to Fab2L allows for the identification of F2 individuals carrying two copies of the transgene from the naive parent (grey chromosomes) rather than from the epiline parent (blue chromosomes). See also **Figure S4**. **(B-F)** Paramutation crossing schemes and phenotypic distribution of the populations with the indicated genotypes and epiline identities. Pie charts represent the phenotypic distribution of the eye colour within the population sorted into five classes.

To determine if mutation of the transgene interfered with horizontal transfer of epigenetic state, we crossed the Fab2L-INS, Fab2L-PRE and Fab2L-INS-PRE lines with a white-eyed Fab2L epiline in order to see if these mutant versions of the Fab2L transgene could acquire a repressed epigenetic state by paramutation in addition to, or instead of, selection over generations. Again, as control we used a wild-type Fab2L epiline which had previously acquired a de-repressed, red-eyed epigenetic state. Just as with the selection, Fab2L, Fab2L-INS and Fab2L-PRE were able to acquire a more white-eyed phenotype than the naïve parental lines (Figure 3C-E). Conversely, all Fab2L-INS-PRE individuals maintained their uniform red coloration of the eyes even after exposure in the F1 of the cross to a repressed epiallele (Figure 3F). While lines bearing a wild-type version of the insulator or PRE of *Fab-7* alone were therefore able to acquire an altered epigenetic state by both selection over generations and paramutation, mutation of both regions together completely prevents acquisition of a repressed epigenetic state by either method. Taken together, these results therefore suggest that the insulator and PRE work together not only to mediate epigenetic variation at the transgenic *Fab-7*, but also to maintain an epigenetic memory across generations.

### GAF mediates long-range chromatin contacts through the insulator region of *Fab-7*

Our results point to PRC2 and GAF as key factors mediating the epigenetic variability, and its inheritance across generations, at the Fab2L transgene. PRC2 has a direct role in regulating the expression of the Fab2L transgene, as the differences in *mini-white* expression and phenotype of the Fab2L epilines correlate with differences in PRC2-deposited H3K27me3 across the transgene^19^. The role of GAF is less clear as mutation of the GAF sites in the insulator ultimately had little effect on PRC2 binding (Figure 1E), suggesting that GAF’s primary role at the *Fab-7* element may not be the recruitment of PRC2. Intriguingly, among its many other functions, GAF has been shown to mediate long-range chromatin contacts between its target genes^31–33^. The three-dimensional organization of chromatin is a major factor in the regulation of gene expression, and polycomb target genes in particular are frequently found to colocalize in the nucleus at so-called “Polycomb bodies”^34^. Moreover, the transgenic and endogenous copies of *Fab-7* were previously found to form chromatin contacts in the Fab2L line^19^.

To quantify these chromatin contacts, we performed Fluorescence in situ hybridisation (FISH) to visualise the regions surrounding the *Fab-7* elements in Fab2L embryos carrying different copy numbers of the endogenous *Fab-7* (Figure 4A,B). These regions showed significant colocalization in the nuclei of Fab2L embryos, but not in Fab2L ; *Fab7[1]* embryos which lack the endogenous *Fab-7*. This shows that chromatin contacts do occur between these loci dependent on the presence of both *Fab-7* elements. Intriguingly, the Fab2L-*Fab-7* chromatin contacts observed in Fab2L increase even further in a *Fab7[1]* / + genetic background in which only one copy of the endogenous *Fab-7* is present.

**Figure 4.**
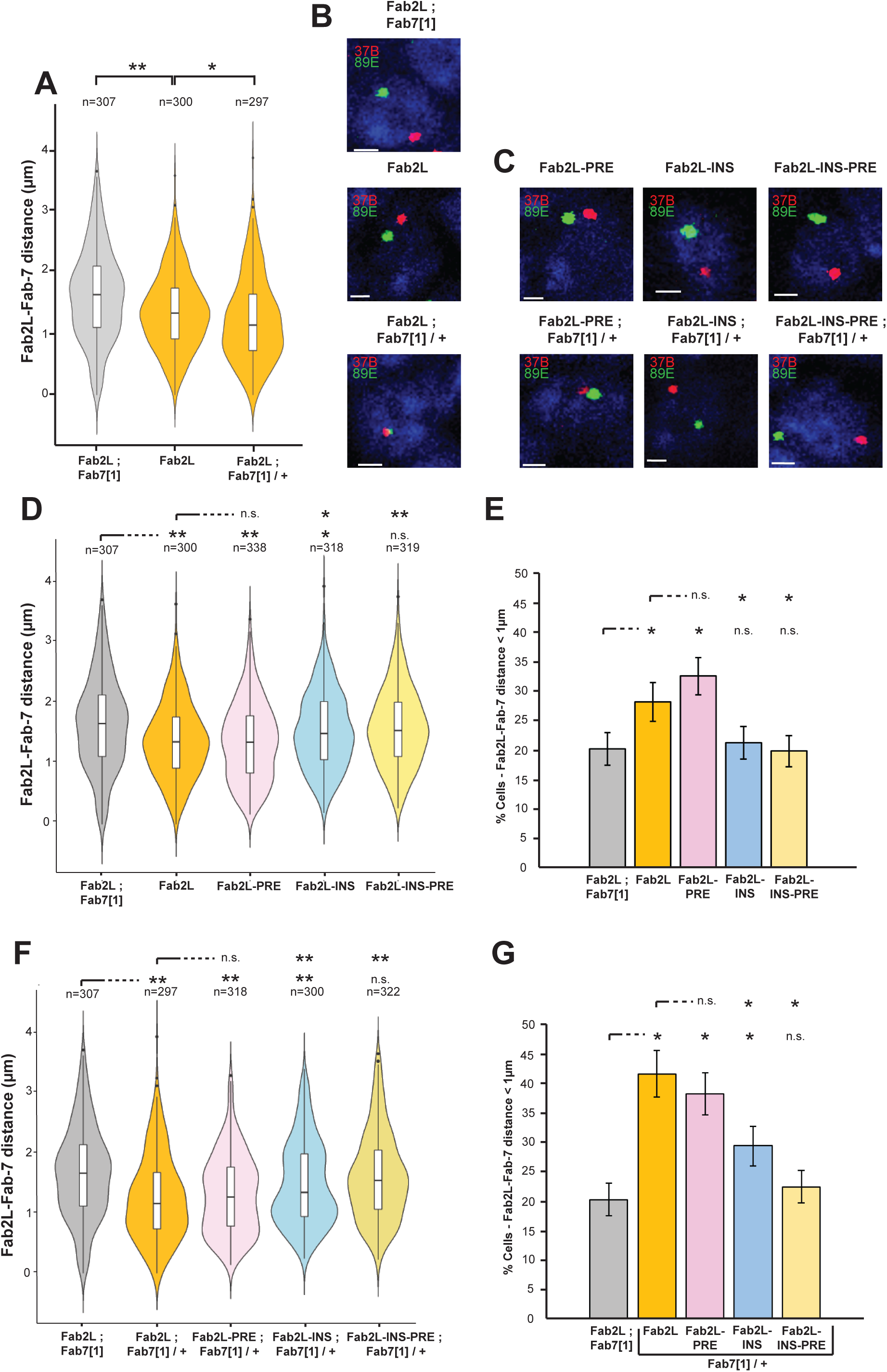
Long-distance chromatin contacts between Fab2L and *Fab-7* depend on the functionality of the insulator region. **(A,D,F)** Violin plots representing the distribution of average distance between the 37B and 89E regions surrounding the Fab2L transgene and endogenous *Fab-7*, respectively, as determined by FISH in the indicated genotypes. Distances were measured in stage 14-15 embryos in T1 and T2 segments. Distributions were compared using the t-test (*p<0.05, **p<0.01, n.s. = not significant). **(B,C)** Illustrative micrographs of FISH in embryonic nuclei of the indicated genotypes. Nuclei are stained with DAPI in blue, the 37B locus surrounding the Fab2L transgene is stained in red and the 89E locus surrounding the endogenous *Fab-7* is stained in green. Scale bars represent 1 μm. See also **Figure S5**. **(E,G)** Bar graphs representing the percentage of cells from the same FISH assays in which the inter-loci distance was less than 1 μm.

To investigate the effect of Fab2L mutation on these chromatin contacts, we extended the FISH analysis to our mutated transgenic lines (Figures 4C-G and S5). In both a homozygous and hemizygous *Fab-7* background, chromatin contacts between Fab2L and *Fab-7* were comparable between Fab2L-PRE and wild-type Fab2L. This is measured both in a similar average distance between the two loci (Figure 4D,F) and in the proportion of nuclei in which the loci are in close proximity (Figure 4E,G). Conversely, contacts are significantly decreased in Fab2L-INS compared to wild-type Fab2L, suggesting a primary role for the insulator, and GAF, in mediating these chromatin contacts. It is worth noting, however, that mutation of the insulator does not fully abrogate chromatin contacts to the extent seen in Fab2L ; *Fab7[1]*, whereas mutation of both the insulator and PRE does. Thus, while Fab2L-PRE is not significantly different from Fab2L, and Fab2L-INS-PRE is not significantly different from Fab2L ; *Fab7[1]*, Fab2L-INS displays an intermediate phenotype, indicating significant, but not complete, loss of contacts between the two loci (Figure 4D-G, and S5C,G). Taken together, these results strongly suggest that contacts between the *Fab-7* elements are primarily mediated through the insulator region, but that the PRE may also play a stabilizing role.

### Artificially-induced chromatin contacts are sufficient to induce TEI at the Fab2L locus

We then sought to investigate whether chromatin contacts may play a causal role in the establishment of TEI at Fab2L. Inter-loci Fab2L-*Fab-7* distance does not differ significantly between white- or red-eyed epilines of Fab2L^19^, arguing against a contribution of chromatin contacts to the maintenance of epigenetic differences between these epilines. We also found that while these contacts are robust in late-stage embryos (stage 14-15) they are not observable in early stages (stage 4-5) (Figure S6), indicating that chromatin contacts themselves are unlikely to be the transgenerationally inherited signal of TEI. However, the increase in Fab2L-*Fab-7* chromatin contacts in Fab2L ; *Fab7[1]*/+ individuals hemizygous for the endogenous *Fab-7* (Figure 4A), correlating with the triggering of TEI in the genetic crosses previously discussed (Figure S2), makes the gain of chromatin contacts a prime candidate for the molecular trigger that establishes TEI. We therefore wished to explore this correlation in greater detail.

To directly investigate the role of chromatin contacts in the triggering of TEI, we therefore sought to induce contacts rather than abrogate them. To achieve this, we developed an *in vivo* system to induce interchromosomal contacts between the two regions of interest without recourse to any genetic perturbation. We dubbed this system “Three-Dimensional Contact Induction System” or “3D-CIS”, for its ability to bring two distant loci in proximity (Figures 5A and S7). To create this system, we inserted arrays of Lac or Tet operators adjacent to the transgenic and endogenous *Fab-7* elements, respectively. Aside from the addition of these arrays, this line has the same genotype as the Fab2L line, and has a similar average distance and contact frequency between the two loci (Figures 5B,D,E and S8A). However, activation of the system by introducing a TetR-LacI fusion protein which binds to both arrays (Figure S7E,F) results in anchoring of the two *Fab-7* elements to each other (Figure 5C). This anchoring leads to a decrease in the average Fab2L-*Fab-7* distance and increase in the frequency of close contacts, both to levels comparable to the hemizygous Fab2L ; *Fab7[1]* / + in which TEI is established (Figures 5D,E and S8B). Importantly, at no point are either the transgenic or endogenous *Fab-7* in a hemizygous state (Figure S7A,B). The 3D-CIS system therefore allows us to investigate the effect of increasing chromatin contacts between the two *Fab-7* elements in the absence of any genetic perturbation.

**Figure 5.**
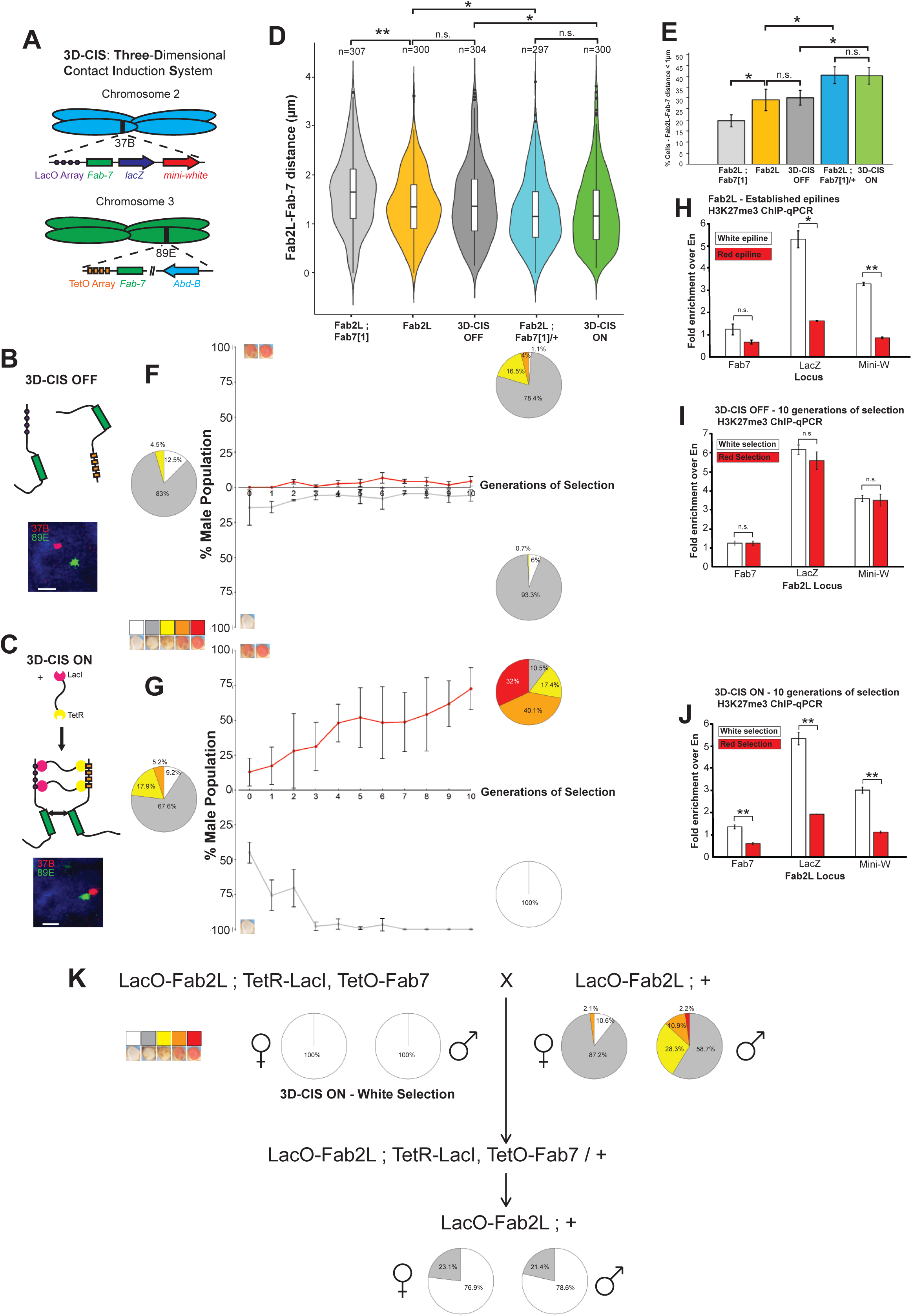
An *in vivo* synthetic biology system promotes interchromosomal contacts between *Fab-7* elements and is sufficient to induce TEI without genetic perturbation. **(A-C)** Schematic representation of the 3D-CIS system. Arrays of Lac and Tet operons are inserted next to the transgenic and endogenous *Fab-7* elements. Expression of a TetR-LacI fusion protein binding to both arrays promotes contacts between the two loci. Illustrative micrographs represent nuclei from embryos with the 3D-CIS system in the “OFF” or “ON” state with FISH highlighting the regions surrounding the *Fab-7* elements. Nuclei are stained with DAPI in blue, the 37B locus is stained in red and the 89E locus is stained in green. Scale bars represent 1 μm. See also **Figures S7 and S8**. **(D)** Violin plots representing the distance distributions of the 37B and 89E regions surrounding the two *Fab-7* elements as determined by FISH in the indicated genotypes. Distances were measured in stage 14-15 embryos in T1 and T2 segments. Distributions were compared using the t-test (*p<0.05, **p<0.01, n.s. = not significant. **(E)** Bar graphs representing the percentage of cells from the same FISH assays in which the inter-loci distance was less than 1 μm. **(F,G)** Results of selection for the most white or red-eyed flies in each generation of 3D-CIS flies in either the “OFF” or “ON” state. Curves represent the percentage of males of Class 1 or Class 4+5 in the population across generations. Error bars are +/- SD of 3 independent repeats. Pie charts represent the phenotypic distribution of the eye colour within the population in the first and last generations of selection, sorted into five classes. See also **Figures S9, S10**. **(H-J)** ChIP-qPCR assays against H3K27me3 performed in embryos of the indicated genotypes after at least 10 generations of selection towards white or red epiallele identity, at regions within the Fab2L transgene. Error bars represent +/- SEM of three independent repeats. Samples were normalised to engrailed as a positive control and compared to each other by the t-test (*p<0.05, **p<0.01, n.s. = not significant). **(K)** Paramutation crossing scheme and phenotypic distribution of the populations with the indicated genotypes and epiline identities. See also **Figure S11**.

Just as with Fab2L in the absence of genetic perturbation (Figure S2A,B), selection of the 3D-CIS line in the “OFF” state over several generations did not result in any change in eye colour across the population, towards either white or red eyes (Figure 5B,F). After ten generations of selection, these lines also exhibited no difference in H3K27me3 levels between each other (Figure 5I). However, activation of the 3D-CIS system, by introduction of the TetR-LacI fusion protein and thus increase in contacts between the Fab2L transgene and endogenous *Fab-7*, was able to establish TEI, such that selection over subsequent generations resulted in both white and red epilines (Figure 5C,G). These epilines also had significant differences in H3K27me3 levels between them (Figure 5J), reminiscent of the differences between Fab2L epilines obtained by selection after transient hemizygosity of *Fab-7* (Figure 5H). These results demonstrate that chromatin contacts alone, in the absence of any genetic perturbation, are sufficient to induce TEI at the Fab2L transgene. As further controls, we generated two more lines expressing either a LacI-LacI or a TetR-TetR fusion protein as part of the 3D-CIS system (Figure S7C,D). These lines were also unable to trigger TEI (Figure S9), showing that triggering is not due to expression of a fusion protein or its binding to either array singly, but conclusively results from the binding of the fusion protein to both arrays in tandem.

### The *Fab-7* element is required for stable chromatin contacts

The ability of the 3D-CIS system to induce TEI in Fab2L suggests that the primary role of the *Fab-7* element in the establishment of TEI is to mediate long-range chromatin contacts between the transgenic and endogenous *Fab-7* elements. To determine whether induced chromatin contacts can trigger TEI at the Fab2L transgene even in the absence of the *Fab-7* element, we generated a new version of the 3D-CIS system in which the transgenic *Fab-7* was deleted (Figure S10A). As expected, phenotypically, this line resembled Fab2L-INS-PRE, with all individuals possessing uniform red eyes (Figures 1J and S10B). Similarly, this LacO-Fab2L-Fab7Δ line was unable to acquire a repressed epigenetic state by either selection or paramutation (Figure S10C-E). Activation of the 3D-CIS system by introduction of the LacI-TetR fusion protein was also unable to trigger TEI in this line (Figure S10F-I). However, FISH analysis revealed that chromatin contacts between the transgene and the endogenous *Fab-7* were not increased in this line. Indeed in both the “OFF” and “ON” state, 3D-CIS-Fab7Δ flies did not show any significant contacts between the two loci, comparable to Fab2L ; *Fab7[1]* (Figure S10J). This suggests that 3D-CIS is insufficient to mediate long-range chromatin contacts on its own, but rather acts to stabilise or reinforce contacts already established between the two *Fab-7* elements.

### Altered epigenetic states remain stable in the absence of artificially-induced chromatin contacts

These results demonstrate a clear role for chromatin contacts in the initial triggering of TEI in Fab2L. However, due to experimental constraints, the 3D-CIS system remains active throughout the selection towards epilines. To determine whether the altered epigenetic states triggered by the 3D-CIS system can be maintained even in the absence of induced chromatin contacts, we crossed the 3D-CIS epilines with a naïve LacO-Fab2L. In the F2, flies lacking the TetR-LacI fusion protein (as determined by a GFP marker, see Methods) were selected and counted. Even in the absence of the TetR-LacI inducing chromatin contacts, this F2 generation had a majority of individuals with primarily white eyes, in the case of the white epiline (Figure 5K), or primarily red eyes, in the case of the red epilines (Figure S11A). The distribution of the population was comparable with those derived from a cross with Fab2L epilines triggered by transient hemizygosity rather than 3D-CIS (Figure S11B,C). The LacO-Fab2L was therefore able to maintain the memory of its altered epigenetic state, even in the absence of artificially-induced chromatin contacts with the endogenous *Fab-7*, demonstrating that enhancement of chromatin contacts is required to establish, but not maintain, transgenerational epigenetic inheritance.

## Discussion

### GAF-mediated chromatin contacts and PRC2-mediated epigenetic variability together account for TEI at the Fab2L transgene

Our results highlight the crucial role played by two subdomains of the *Fab-7* element in the establishment of epigenetic variation at the Fab2L transgene, and the maintenance of its memory across generations. These subdomains are an insulator and a PRE, which act through the recruitment of the transcription factors GAF and Pho. Alone, one of these regions remains sufficient to maintain a certain degree of variation and epigenetic memory at the Fab2L transgene, albeit in a manner skewed towards de-repression. Mutation of all GAF and Pho sites across both regions, however, completely abrogates all variation, demonstrating that at least some binding of these proteins is essential (Figures 1-2).

Our findings suggest that the insulator and PRE cooperate to control epigenetic regulation of the *Fab-7* element in two ways. The first is the recruitment of PRC2, which deposits H3K27me3 in a stochastic manner, leading to the observed variable eye colour phenotype. The second is to mediate long-range chromatin contacts between the two distant *Fab-7* elements in the genome. However, mutation of these subdomains suggests that the PRE is the comparatively more important of the two regions for PRC2 recruitment (and thus epigenetic variability) (Figure 1E), while the insulator is much more involved in the establishment of chromatin contacts (Figure 4). Nevertheless, both elements contribute to some extent to both aspects of Fab2L regulation.

The distribution of GAF and Pho bindings sites between these subdomains suggests differing roles for these proteins, in agreement with what is known of their function. Indeed, the majority of GAF binding sites (6 out of 9) are located in the insulator, while the majority of Pho sites (3 out of 4) are in the PRE (Figure 1B). Based on these observations, we propose that GAF is the primary mediator of chromatin contacts, whereas Pho is the primary recruiter of PRC2, and thus responsible for the epigenetic variability at the Fab2L transgene. Together, these proteins thus mediate the dual functions of the *Fab-7* element, both of which are essential to TEI in this model system.

### PRC2-dependent epigenetic memory at the Fab2L locus

Our results clearly point to a central role for chromatin contacts in the establishment of TEI at Fab2L, but not in the inheritance of the alternative gene expression at Fab2L, as contacts are not present in early development (Figure S6). PRC2-deposited H3K27me3 is thus the primary epigenetic signal underpinning the variability at the Fab2L transgene and must be inherited by another mechanism independent of chromatin contacts, either by direct inheritance or by reconstruction in each generation based on an epigenetic memory maintained by some other signal^6^.

Alternatively, it is worth considering that a combination of direct and indirect inheritance mechanisms could serve to reinforce each other, providing a more stable epigenetic memory that either pathway alone. Previous work in *Drosophila* has provided evidence of germline inheritance of H3K27me3, arguing for its ability to be transmitted through gametogenesis at least^35^. Moreover, a recent study found that a DNA-binding protein, in this case CTCF, can remain bound to chromatin through development and maintain an epigenetic memory through this association^20^. That this mechanism could extend to a complex like PRC2, which both binds to and deposits H3K27me3, is an interesting prospect, as it has the potential to provide a positive feedback loop to stabilise transgenerational H3K27me3. In this way, inheritance of H3K27me3 both directly, by transmission through the germline, and indirectly, through a memory of PRC2 binding, could provide redundancy and reinforcement to ensure a more reliable inheritance of epigenetic memory. While our study is primarily concerned with explaining how TEI is triggered at the Fab2L locus, future investigations into the mechanism by which this epigenetic state is transmitted across generations will provide a more complete picture of this instance of TEI.

### Chromatin contacts trigger PRC2-dependent TEI at the Fab2L locus

Our results indicate that the primary mechanism for the triggering of TEI at the Fab2L transgene is the promotion of physical contact within the nucleus between Fab2L and the endogenous *Fab-7*. Increased contact frequency can either be induced by hemizygosity (Figure S2) – probably because the remaining copy of *Fab-7* forms contacts more efficiently with its distant homolog at another locus in the absence of its homologous allele – or in a synthetic manner, such as in our transgenic 3D-CIS system (Figure 5). This demonstrates that chromatin contacts can establish transgenerational epigenetic memory in the absence of any genetic perturbation. Nevertheless, we note that this system is unable to mimic the effects of hemizygosity in the absence of an adjacent *Fab-7* element (Figure S10). This is explained by the fact that Fab2L-*Fab-7* contacts are already observed to a lesser extent in Fab2L ; + individuals, but are increased in Fab2L ; *Fab[7]*/+ hemizygotes (Figure 4A). Thus, 3D-CIS acts to increase or stabilise the contacts already occurring between the two loci, rather than driving the contacts *de novo*.

As this trigger can lead to inheritance of epigenetic state in both directions (repression and de-repression), any mechanism explaining how this trigger occurs must take into account this plasticity. We therefore propose a model whereby stabilisation of Fab2L-*Fab-7* chromatin contacts allows for the exchange of PRC2 between the endogenous and transgenic *Fab-7* elements, thereby triggering an epigenetic memory that can be selected towards extremes over generations (Figure 6). In this model, naïve Fab2L flies have stochastic recruitment of PRC2 to the transgene by Pho, leading to a random mosaic eye colour pattern (Figure 6A). Chromatin contacts are mediated by GAF, but in the absence of manipulation these contacts primarily occur between homologous alleles, i.e. Fab2L to Fab2L or endogenous *Fab-7* to endogenous *Fab-7* (Figure 6B). While interchromosomal contacts between Fab2L and *Fab-7* do occur, they are transient and outcompeted by the preferential interaction between homologous alleles that is common in dipteran species^36^ (Figure 6C). Stabilisation of these contacts is achieved upon *Fab-7* hemizygosity (Figure 6D), because the remaining endogenous *Fab-7,* having lost its preferred interaction partner, is free to form more stable contacts with its imperfect transgenic partner without being outcompeted by its homologous allele. This situation can be mimicked in a homozygous state, and in the absence of genetic perturbation, thanks to the 3D-CIS transgenic system, which artificially stimulates chromatin contacts between Fab2L and *Fab-7*, making them interact preferentially with each other rather than with their homologous alleles (Figure 6E).

**Figure 6.**
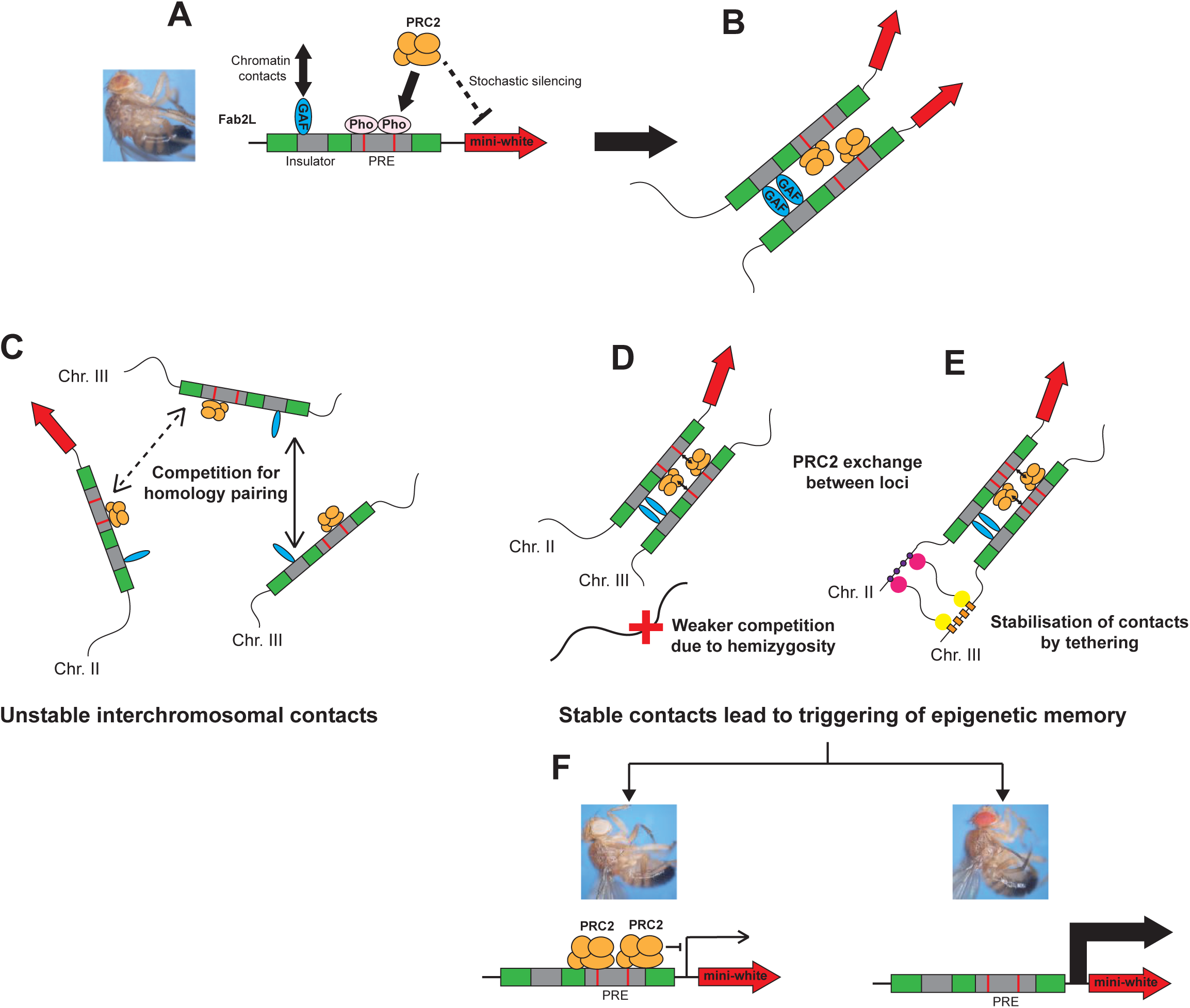
Model: Stabilisation of interchromosomal contacts triggers an epigenetic memory of PRC2 binding. **(A)** The *Fab-7* element recruits PRC2 by Pho binding to its PRE, leading to stochastic silencing of a *mini-white* transgene and a mosaic eye colour. **(B)** This PRC2 recruitment is coupled with long-range chromatin contacts with other *Fab-7* elements mediated by GAF through an insulator region. **(C)** When more than one copy of *Fab-7* is present in the genome contacts can be initiated between distant *Fab-7* elements, but these contacts are outcompeted by inter-allelic contacts and remain transient. (**D-E**) Stabilization of these contacts can be achieved through hemizygosity of one *Fab-7* copy (D) or through synthetic biology tools (E). This stabilisation leads to exchange of PRC2 between the PREs, resulting in either increased PRC2 association and silencing, or decreased PRC2 association and de-repression, and triggering an epigenetic memory of this altered association. **(F)** Over generations these slight differences can be selected to extremes, resulting in either very strong or very weak repression and strikingly different phenotypes.

When these trans interactions are sufficiently stable, exchange of PRC2 can occur between the PREs of the *Fab-7* elements. Pho sites within the PRE can act as either donors or acceptors of PRC2, leading to small changes in expression of the transgenic *mini-white* reporter. The memory of this expression is maintained across generations, meaning that over time, these differences can be selected to extremes, leading to either fully repressed or fully de-repressed transgene expression, and monochrome eye colour (Figure 6F).

A few aspects of this model are worth highlighting. First, it would predict that a Fab2L/+ ; *Fab7[1]*/+ double hemizygote, in which both loci have lost their preferential interaction partner on the homologue, should be an even more effective trigger of TEI than single hemizygosity. While we have not examined this in detail, our different TEI triggering crosses support this, as epilines derived from Fab2L/+ ; *Fab7[1]*/+ double hemizygotes tend to reach fixation faster than those derived from Fab2L ; *Fab7[1]*/+ single hemizygotes during selection (Figure S2). Second, exchange of PRC2 binding between Fab2L alleles, rather than between Fab2L and *Fab-7*, could also explain how paramutation is able to transfer epigenetic state, albeit imperfectly, between homologous alleles in this line. Our model thus accounts for both the triggering of epiallelic identity, and its horizontal transfer by paramutation.

### A broader role for hemizygosity and chromatin contacts in triggering TEI

One major question in the field of epigenetic inheritance is how heritable epigenetic variability, or epimutation, arises in the first place. Studies in plants suggest that heritable changes in DNA methylation can occur apparently spontaneously in these organisms, leading to long-term epigenetic differences between lines^37–39^. Recent studies in *Caenorhabditis elegans* have extended these observations to metazoans and to RNA and chromatin-based epigenetic changes^40,41^. Other studies have sought to identify environmental triggers for TEI, directly linking epigenetic variation to an external stress to which it is intended to respond^2,42,43^. The final prominent candidate for sources of epigenetic variation is genetic perturbation, different types of which have been shown to trigger TEI in a variety of organisms^15,16,44,45^.

In the *Drosophila* Fab2L line, this genetic perturbation takes the form of transient hemizygosity of the endogenous *Fab-7* region for at least one generation. It is interesting to note that unlike some cases of genetically-triggered TEI, in Fab2L the epigenetic memory is triggered and maintained not at the locus which is perturbed (the endogenous *Fab-7*) but elsewhere in the genome (the Fab2L transgene), testifying to the ability of trans-interactions to induce TEI. *Fab-7* has been found to have a similar effect on other loci. Indeed, hemizygosity of *Fab-7* has been shown to affect the expression of another PRE-containing gene in the distant Antennapedia cluster, with a phenotype that persisted for several generations after restoration of *Fab-7* homozygosity^19^. This raises the question of whether similar mechanisms could be acting to trigger TEI at other loci in natural populations.

Recent sequencing of wild *Drosophila melanogaster* lines has revealed the incredible genomic variation between populations of this single species. This includes numerous and large-scale deletions, duplications and translocations across the genome^46^. Mixing of two such genomically disparate populations would lead to a number of hemizygosity or heterozygosity events, as well as homology between very distant loci reminiscent of what is observed in Fab2L. Breeding in the wild thus has the potential to lead to many instances of naturally occurring genetic perturbation as potential triggers for TEI. Just as in Fab2L, it could be that establishment of TEI might be possible only between certain regions which are already prone to contact each other. In this respect, PRC2 targets may be particularly interesting candidates for naturally occurring TEI. Indeed, as previously mentioned, many PRC2 targets are frequently clustered within the nucleus in polycomb bodies, forming a large domain of silenced chromatin^47^. Interestingly, genes regulated in this manner are more likely to possess both an insulator region and a PRE, just like *Fab-7*. Our study provides insight into the mechanisms by which this type of epimutation could occur, but it is only by extending this insight to a broader context that we will be able to determine the role of TEI in the phenotypic variation, and thus potentially adaptation, of natural populations.

## Acknowledgements

We thank Montpellier Resources Imaging facility as well as the *Drosophila* facility (both affiliated to BioCampus University of Montpellier, CNRS, INSERM, Montpellier, France). We thank Yuki Ogiyama for assistance with the cloning of the plasmids for *Drosophila* injection and Judith Kassis for the gift of the anti-Pho antibody. M.F-J. and G.S. were supported by the European Research Council Advanced Grant 3DEpi with additional support for M. F.-J from the MSD Avenir Foundation Grant GENE-IGH. F.B. and G.C. were supported by CNRS. Research in the P.S. lab was supported by the John Fell Fund (Grant No 0011417). Research in the G.C. laboratory was supported by grants from the European Research Council (Advanced Grant 3DEpi), the European CHROMDESIGN ITN project (Marie Skłodowska-Curie grant agreement No 813327), the European E-RARE NEURO DISEASES grant “IMPACT”, by the Agence Nationale de la Recherche (PLASMADIFF3D, grant N. ANR-18-CE15-0010, LIUVCHROM, grant N. ANR-21-CE45-0011), by the Fondation ARC (EpiMM3D), by the MSD Avenir Foundation ((Project GENE-IGH), and by the French National Cancer Institute (INCa, PIT-MM grant N. INCA-PLBIO18-362).

## Author contributions

M. F-J. and G.C. conceived of and led the project. M. F-J. designed and performed the experiments. M. F-J. and G.C. interpreted the data. M. F-J., G.S. and F.B. performed the *Drosophila* transgenerational selection experiments. M. F-J. composed the manuscript with editorial input from G.C. and P.S. All authors reviewed and commented on the manuscript.

## Methods

### Fly stocks and culture

Flies were raised in standard cornmeal yeast extract media. Standard temperature was 21°C, with the exception of P0 and F1 crosses in the experiments of Fab2L epiallele establishment by hemizygosity or 3D-CIS (Figures 2; 5F,G; S2; S7A-D, S9D,E and S10D-I), for which temperature was 18°C. The Fab2L and Fab2L ; *Fab7[1]* lines were described in Bantignies *et al.*, 2003^22^. The Fab2L, *black[1]* line and pre-established Fab2L epilines (Fab2L-R* and Fab2L-W*) were described in Ciabrelli *et al.*, 2017^19^.

For generation of the Fab2L mutant lines, a transgenic line was made containing an AttP insertion site at cytogenetic position 37B, by CRISPR-Cas9 of a w[1118] line (Bloomington Drosophila Stock Center) to cut at the exact site of Fab2L transgene insertion in the Fab2L line. Fab2L-INS was then generated by Phi-recombination of an AttB-containing plasmid containing the entirety of the Fab2L transgene, with directed mutations of the insulator GAF and Pho sites, into this 37B-AttP line. Injection services for these two lines were provided by BestGene Inc.. Fab2L-PRE and Fab2L-INS-PRE were then generated by CRISPR-Cas9 editing of Fab2L-INS, with a two-guide RNA strategy designed and implemented by Rainbowgene Transgenic Flies Inc.

To create the 3D-CIS system, arrays of 7 Tet operators and 21 Lac operators were inserted adjacent to the endogenous or transgenic *Fab-7*, respectively, using a single guide RNA to cut immediately to their 3’. Cassettes encoding recombinant proteins combining the Lac and/or Tet repressors with a GFP marker (TetR-GFP-LacI, TetR-GFP-TetR and LacI-GFP-LacI) under expression of an *Actin-5C* promoter were inserted into chromosome arm 3L separately by Phi recombination into an established AttP containing line (Bloomington 24480). The transgenes encoding these proteins were then recombined with the TetO-*Fab-7*, and introduced into a Fab2L background, ready to be crossed with the LacO-Fab2L as described in Supplementary information, Figure S7. All injections for these lines were provided by BestGene Inc. The Fab2L-*Fab-7*Δ line was derived from the LacO-Fab2L line by CRISPR-Cas9 targeted deletion of the transgenic *Fab-7*, designed and implemented by Rainbowgene Transgenic Flies Inc.

Fab2L-INS, *black[1]*, Fab2L-PRE, *black[1]*, Fab2L-INS-PRE, *black[1]* and Fab2L-Fab7Δ, *black[1]* were generated by recombining the Fab2L transgene with the *black[1]* allele from the w[1118]; *black[1]* line (Bloomington Drosophila Stock Center).

### Chromatin Immunoprecipitation and antibodies

0 to 16 hour old embryos were collected in Embryo Wash Buffer (0.03% Triton X-100, 140mM NaCl) and dechorionated with bleach. Samples were crosslinked in 1 ml A1 buffer (60 mM KCl, 15 mM NaCl, 15 mM HEPES [pH 7.6], 4 mM MgCl2, 0.5% Triton X-100, 0.5 mM dithiothreitol (DTT), 10 mM sodium butyrate and complete EDTA-free protease inhibitor cocktail [Roche]), in the presence of 1.8% formaldehyde. Samples were homogenized with a micropestle and incubated for a total time of 15 minutes at room temperature. Crosslinking was stopped by adding 350 mM glycine followed by incubation for 5 min. The homogenate was transferred to a 2 ml tube and centrifuged for 5 minutes, 4,000g at 4°C. The supernatant was discarded, and the nuclear pellet was washed three times in 2 ml A1 buffer and once in 2 ml of Lysis buffer (140 mM NaCl, 15 mM HEPES [pH 7.6], 1 mM EDTA, 0.5mM EGTA, 1%Triton X-100, 0.5mMDTT, 0.1% sodium deoxycholate, 10 mM sodium butyrate and complete EDTA-free protease inhibitor cocktail [Roche]) at 4°C. Nuclei were than resuspended in 1.5 ml Lysis buffer in the presence of 0.1% SDS and 0.5% N-Laurosylsarcosine, transferred to a 15 ml falcon tube and incubated for 2 hours with agitation at 4°C. Samples were adjusted to 3 ml and chromatin was sonicated using a Q700 sonicator with microtip (QSonica) for a total of 6 minutes and 30 seconds at amplitude 50 (settings: 30 s on, 1min 30 s off x 13 cycles) in an ice bucket. Sheared chromatin had size range of 100 to 300 base pairs. After sonication and 5 minutes high-speed centrifugation at 4°C, fragmented chromatin was recovered in the supernatant and aliquoted in 5 μg (for H3K27me3 ChIP) or 20 μg (for non-histone protein ChIP) aliquots adjusted to a volume of 500 μl in Lysis Buffer with 0.1% SDS and 0.5% N-Laurosylsarcosine for storage at -20°C.

To perform the ChIP, samples were thawed on ice and chromatin was precleared by addition of 15 μl of Protein A Dynabeads (Invitrogen 10002D) followed by incubation for at least 1 hour at 4°C. Dynabeads were removed on a magnetic rack and antibodies were added at a dilution of 1:100 (a mock control in the presence of rabbit IgG was performed at the same time, while an input of the same size was set aside). Samples were incubated for overnight at 4°C on a rotating wheel. 30 μl of Protein A Dynabeads were added and incubation was continued for at least 2 hours at 4°C. Antibody-protein complexes bound to beads were washed 4 times in Lysis Buffer with 0.05% SDS and twice in TE Buffer (0.1 mM EDTA, 10 mM Tris (pH 8)) in 1 ml each time. Chromatin was eluted from beads in 100 μl of 10 mM EDTA, 1% SDS, 50 mM Tris (pH 8) at 65°C for 15 minutes and eluted again in 150 μl of 10 mM EDTA, 0.67% SDS, 50 mM Tris (pH 8) at 65°C for 15 minutes, with the eluate collected on a magnetic rack each time. The 250 μl eluates and 250 μl of the Input DNA samples (1:2 input) were incubated overnight at 65°C to reverse crosslinks and treated with Proteinase K for 3 hours at 56°C. DNA was isolated by addition of an equal volume of phenol-chloroform, supernatants collected and then ethanol precipitated for 2 hours at -20°C in the presence of 20 μg glycogen by addition of 25 μl 3M sodium acetate and 625 μl ethanol. Samples were centrifuged at high speed for 1 hour and washed in 500 μl of 70% ethanol before resuspension in 200 μl H2O. Immunoprecipitated DNA was used to analyze the enrichment of specific DNA fragments by real-time PCR (qPCR), using a Roche Light Cycler 480 and the Light Cycler 480 SYBR green I Master mix. For each amplicon, IP DNA was normalized to Input DNA. The ChIP/Input ratio was further normalized to a positive control region (engrailed). ChIP amplicons for the insulator or PRE regions were specific to either the WT or mutated transgenic sequence, depending on the genotype analysed. Antibodies used in this study were as follows: anti-GAF polyclonal antibody^48^; anti-Pho polyclonal antibody^48,49^; anti-E(z) polyclonal antibody^31^; anti-H3K27me3 polyclonal antibody (Active Motif 39155), anti-GFP polyclonal antibody (Abcam ab290), normal rabbit IgG (Cell Signalling 2729).

### Fluorescence in situ hybridization

Two-color 3D FISH was performed as previously described^50^. For a detailed protocol, see Bantignies and Cavalli, 2014^51^. Briefly, embryos were dechorionated with bleach and fixed in buffer A (60 mM KCl; 15 mM NaCl; 0.5 mM spermidine; 0.15 mM spermine; 2 mM EDTA; 0.5 mM EGTA; 15 mM PIPES, pH 7.4) with 4% paraformaldehyde for 25 min in the presence of heptane. Embryos were then devitellinized by adding methanol to the heptane phase, extracted and washed three times in methanol. Embryos were kept for a maximum of 4 months in methanol at 4C before proceeding to FISH. Fixed embryos were sequentially re-hydrated in PBT (PBS, 0.1% Tween 20) before being treated with 100–200 μg/ml RNaseA in PBT for 2 hours at room temperature. Embryos were then sequentially transferred into a pre-Hybridization Mixture (pHM: 50% formamide; 4XSSC; 100 mM NaH2PO4, pH 7.0; 0.1% Tween 20). Embryonic DNA was denatured in pHM at 80°C for 15 minutes. The pHM was removed, and denatured probes diluted in the FISH Hybridization Buffer (FHB: 10% dextransulfat; 50% deionized formamide; 2XSSC; 0.5 mg/ml Salmon Sperm DNA) were added to the tissues without prior cooling. Hybridization was performed at 37°C overnight with gentle agitation. Post-hybridization washes were performed, starting with 50% formamide, 2XSSC, 0.3% CHAPS and sequentially returning to PBT. After an additional wash in PBS-Tr, DNA was counterstained with DAPI (at a final concentration of 0.1 ng/μl) in PBT and embryos were mounted with ProLong Gold Antifade (Invitrogen).

FISH probes for the 37B and 89E regions were made from a previous design described in Ciabrelli et al. 2017^19^. For each region, 6 non-overlapping probes of between 1.2 and 1.7kb covering an area of approximately 12kb were generated using the FISH Tag DNA kit with Alexa Fluor 555 or Alexa Fluor 647 dyes (Invitrogen Life Technologies). 100ng of each probe were added to the 30µL of FHB for hybridization.

### Microscopy and image analysis

For the FISH, the 3D distances between 37B and 89E loci were acquired and measured as follows: due to somatic pairing of homologous chromosomes in *Drosophila*, the majority of the nuclei in embryos show a single FISH spot for each probe. In the cases of non-overlap FISH signals between homologues, the closest distance between the centres of the two probes was considered. To measure distances, 3D stacks were collected from 3-5 different embryos. Optical sections were collected at 0.5 μm intervals along Z-axis using a Leica SP8-UV microscope, Montpellier Resources Imaging (MRI) facility. Relative 3D distances between FISH signals were analyzed in approximately 80 to 120 nuclei per 3D stack using the Imaris software (Oxford Instruments). The distance distribution between the two probes was obtained by pooling replicates for each condition.

## Supplemental Information Titles & Legends

**Figure S1.**
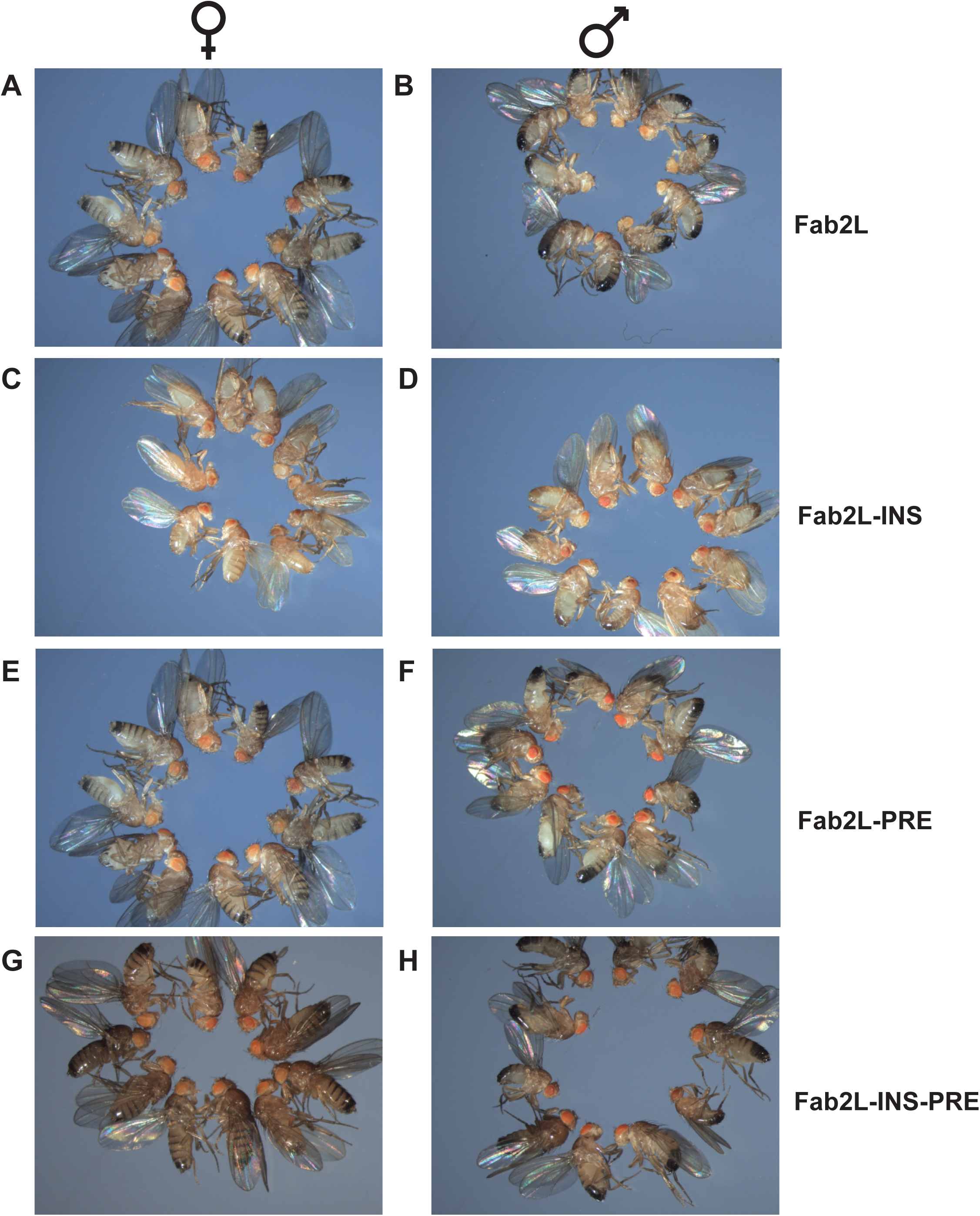
Mutation of GAF and Pho sites leads to stronger red eye colour phenotypes. **(A-H)** Images showing randomly selected samples of 10 females and males of the indicated genotypes.

**Figure S2.**
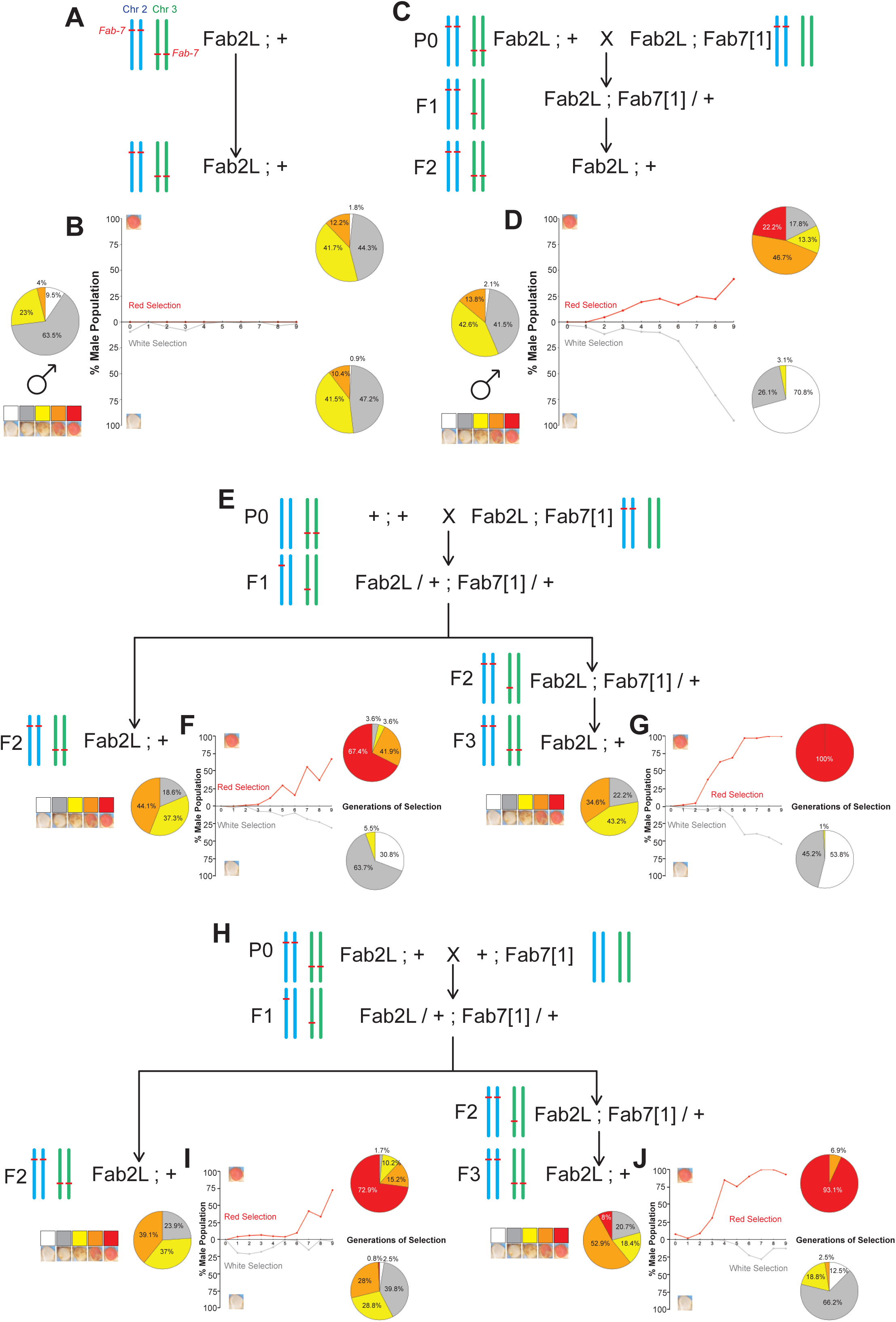
Transient hemizygosity of *Fab-7* triggers transgenerational epigenetic inheritance at the Fab2L transgene. **(A-J)** Crossing schemes and results of subsequent selection towards epilines of the Fab2L transgene. Selection was performed after either no cross (A,B), transient hemizygosity of the endogenous *Fab-7* element (C,D), or transient hemizygosity of both the endogenous and transgenic *Fab-7* elements, achieved from crosses with different parental genotypes (E-J). In each case, Fab2L ; + flies derived from the indicated cross were split into two independent lines and subjected to repeated selection of the most extreme white or red-eyed flies, respectively, in each generation. Curves represent the percentage of Class 1 or Class 5 males in the population across generations. Pie charts represent the phenotypic distribution of the eye colour within the population in the first and last generations of selection.

**Figure S3.**
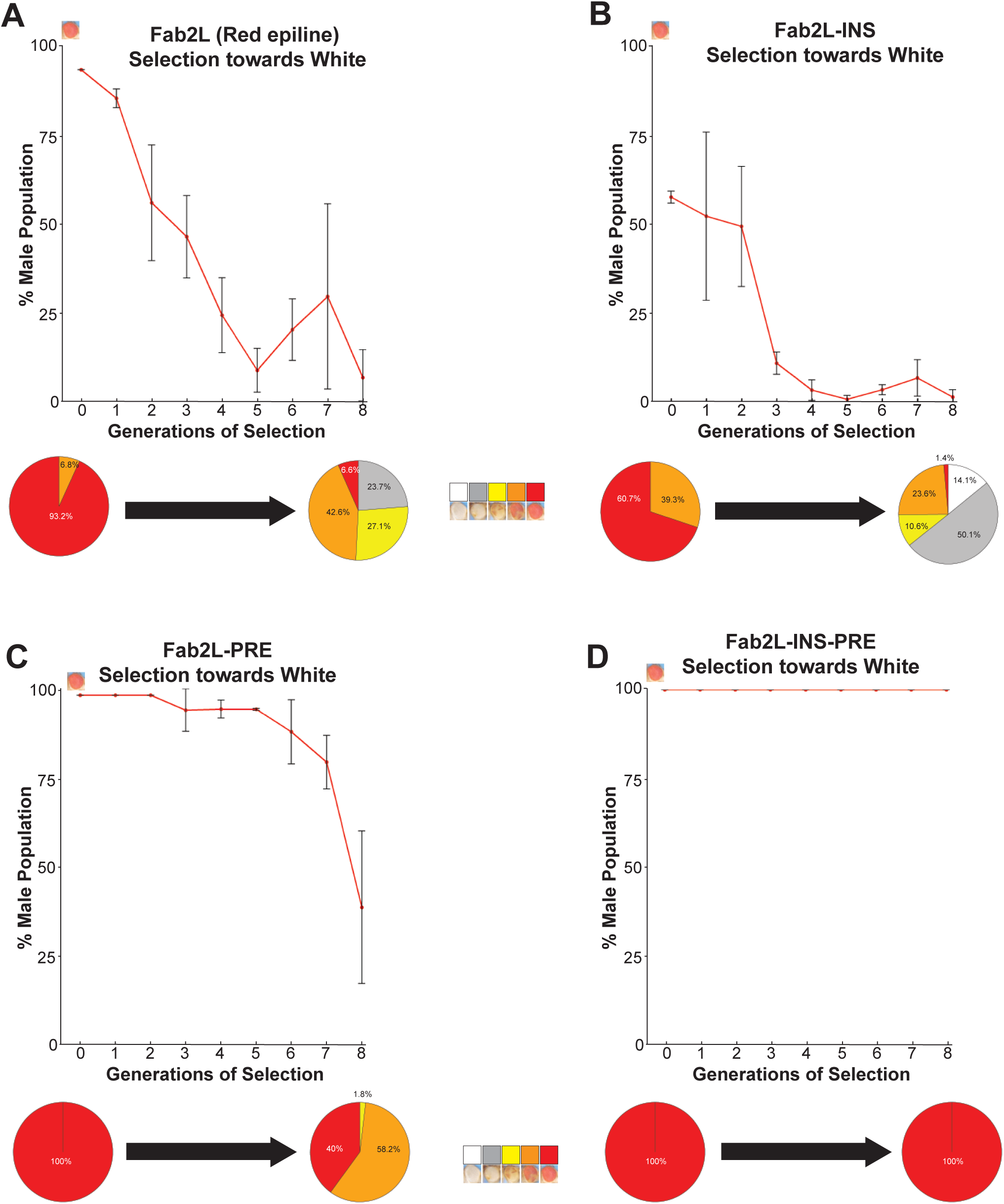
Epigenetic inheritance in males is abrogated in the absence of GAF and Pho binding to the Fab2L transgene. **(A-D)** Results of selection for the most white-eyed flies in each generation beyond the F2 of the crossing scheme. At top, curves represent the percentage of Class 5 males in the population across generations. Error bars are +/- standard deviation (SD) of 3 independent repeats. At bottom, pie charts represent the phenotypic distribution of the eye colour within the population in the first and last generations of selection, separated into five classes.

**Figure S4.**
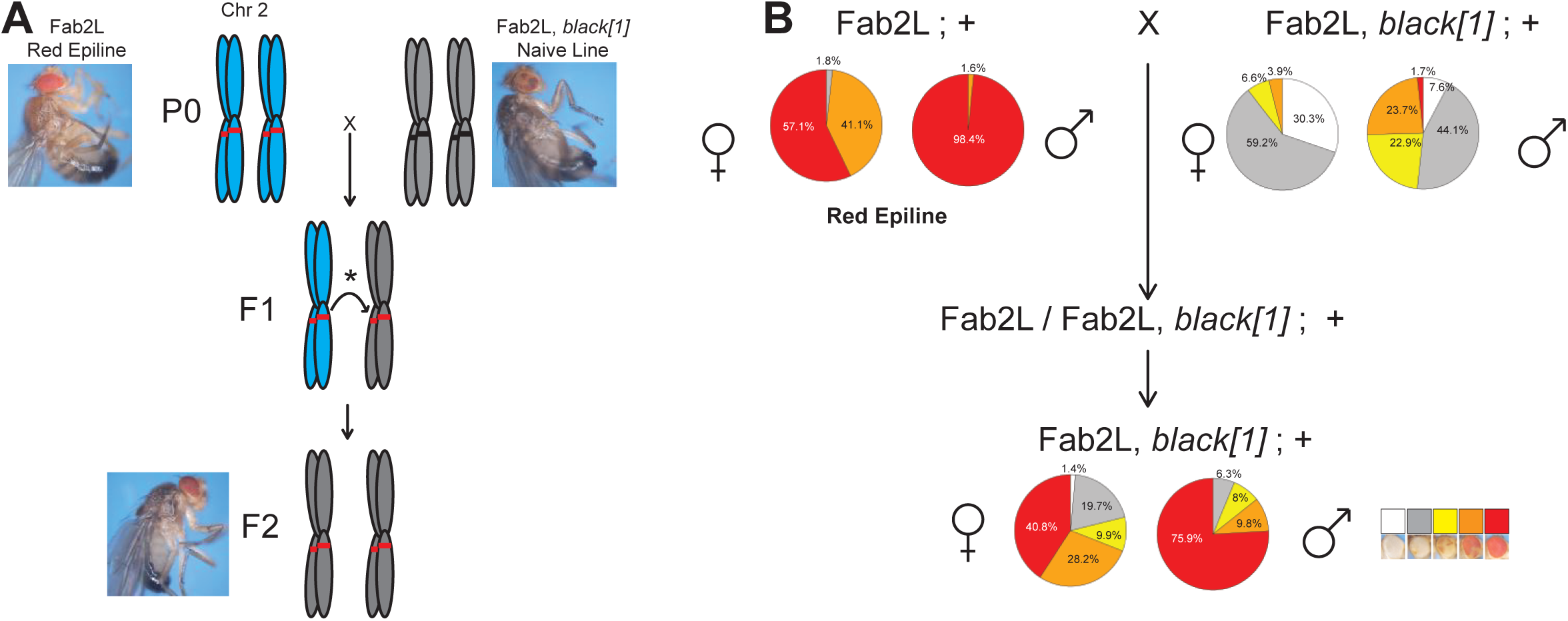
Fab2L can acquire a derepressed epigenetic state *in trans* by paramutation. **(A)** Illustration of the paramutation crossing scheme for the acquisition of a derepressed epigenetic state by a naive Fab2L allele from an established epiallele *in trans*. **(B)** Paramutation crossing scheme and phenotypic distribution of the populations with the indicated genotypes and epiline identities. Pie charts represent the phenotypic distribution of the eye colour within the population separated into five classes.

**Figure S5.**
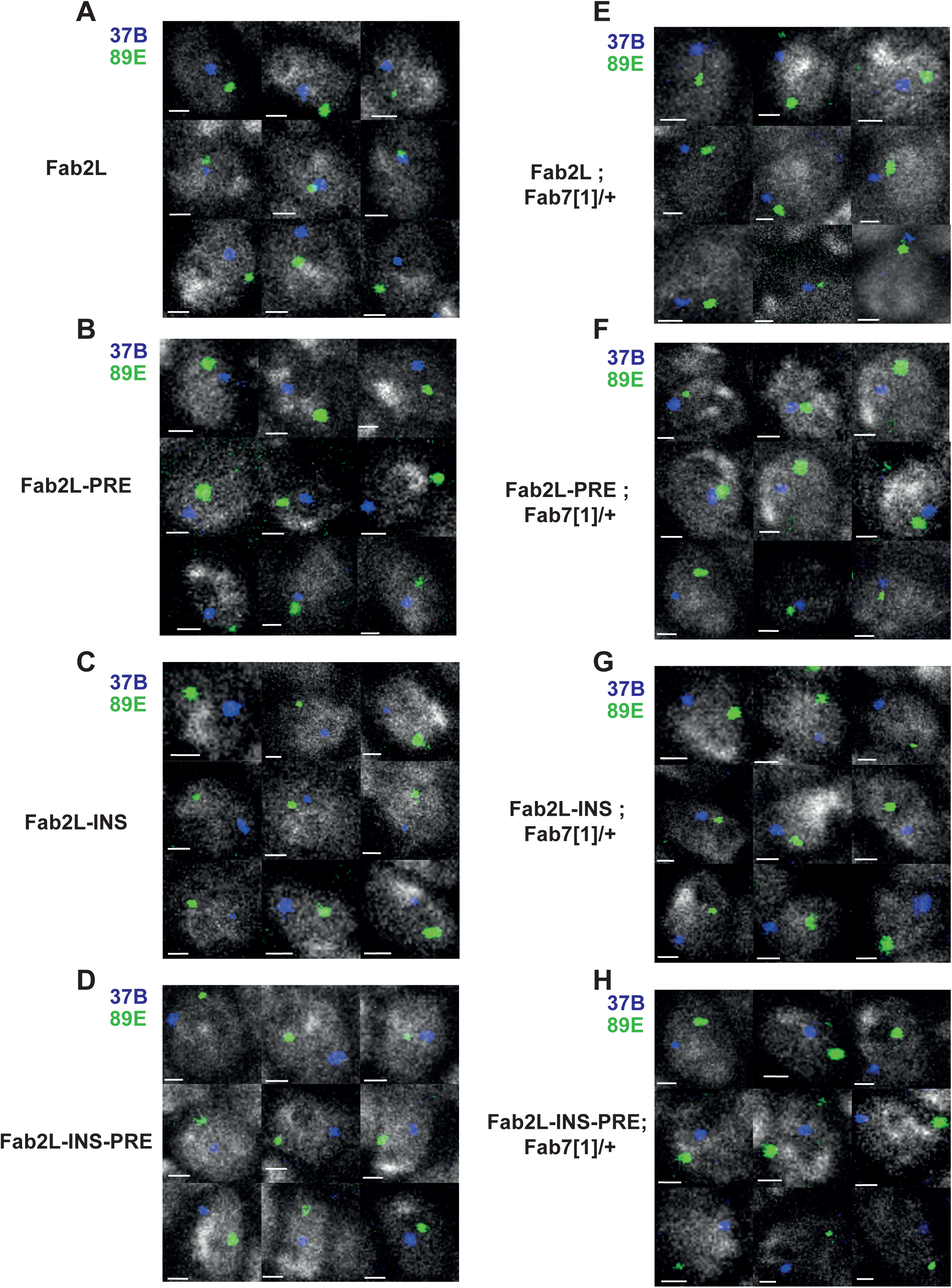
Long-distance chromatin contacts between Fab2L and *Fab-7* increase upon *Fab-7* hemizygosity, but not in insulator mutants. **(A-H)** Micrograph galleries of randomly selected nuclei of the indicated genotypes in FISH-stained embryos. Nuclei are stained with DAPI in grey, the 37B locus surrounding the Fab2L transgene is stained in blue and the 89E locus surrounding the endogenous *Fab-7* is stained in green. Scale bar represents 1 μm. The quantification associated with these images is represented in Figure 4.

**Figure S6.**
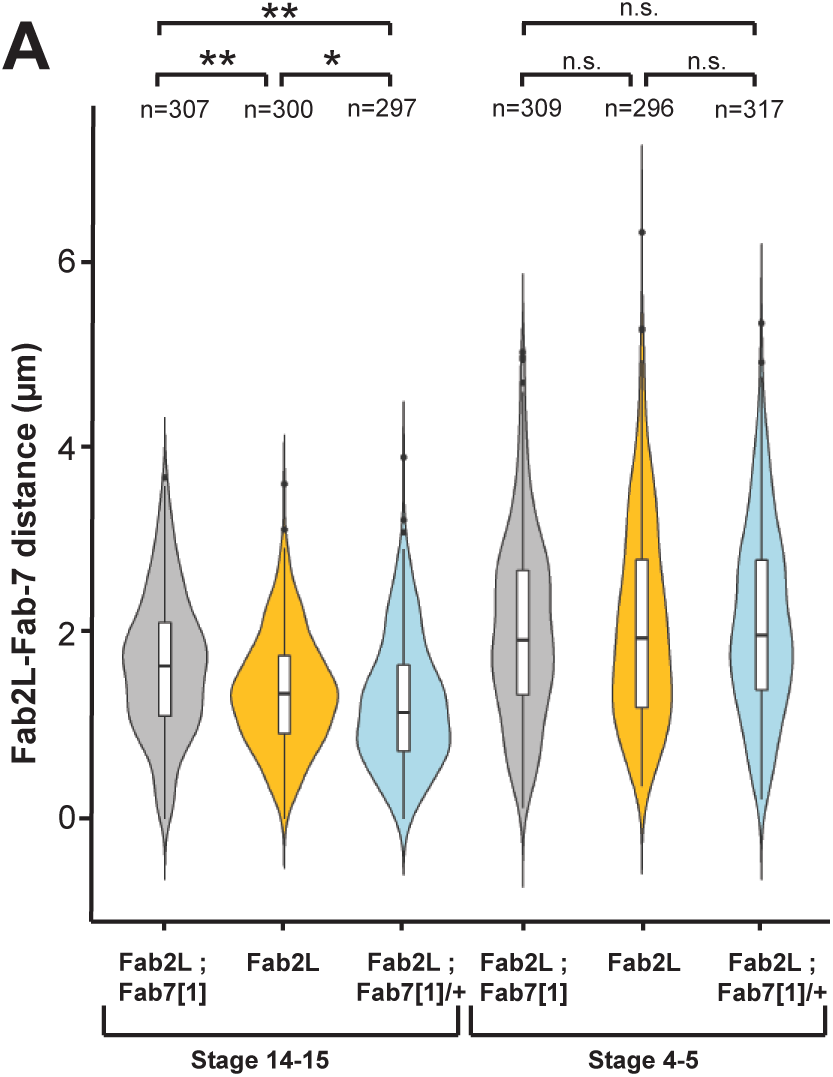
Chromatin contacts between *Fab-7* elements are not present in the early embryo. **(A)** Violin plots representing the distance distributions of the 37B and 89E regions surrounding the two *Fab-7* elements as determined by FISH in the indicated genotypes and embryo stages. Distributions were compared using the t-test (*p<0.05, **p<0.01, n.s. = not significant).

**Figure S7.**
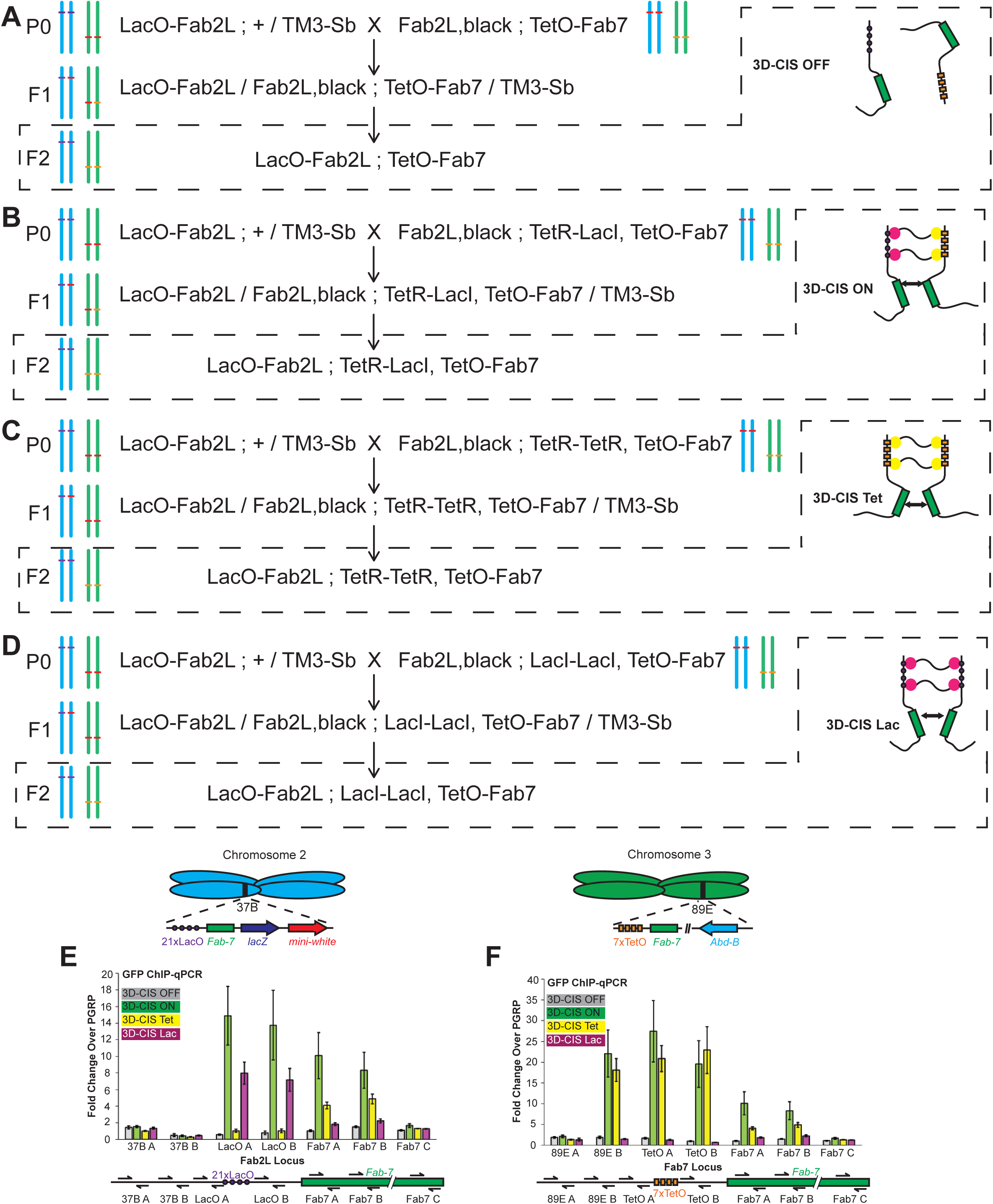
Synthetic constructs of the 3D-CIS system and its variants bind to their target sequences. **(A-D)** Crossing schemes used to generate the 3D-CIS lines and associated controls. Chromosomes 2 and 3 for each genotype of the crosses are illustrated with the different versions of the *Fab-7* element highlighted. Red bars indicate a wild-type *Fab-7* at either the endogenous (chromosome 3) or transgenic Fab2L (chromosome 2) locus. Purple and orange bars indicate the array-containing LacO-Fab2L and TetO-Fab7, respectively. As illustrated, none of the crosses introduce hemizygosity of either *Fab-7* element at any point. **(E,F)** ChIP-qPCR assays in embryos of the indicated genotypes against LacI-TetR, TetR-TetR or LacI-LacI using a GFP tag. Approximate positions of the loci analysed by each primer pair within or adjacent to the *Fab-7* elements are indicated.

**Figure S8.**
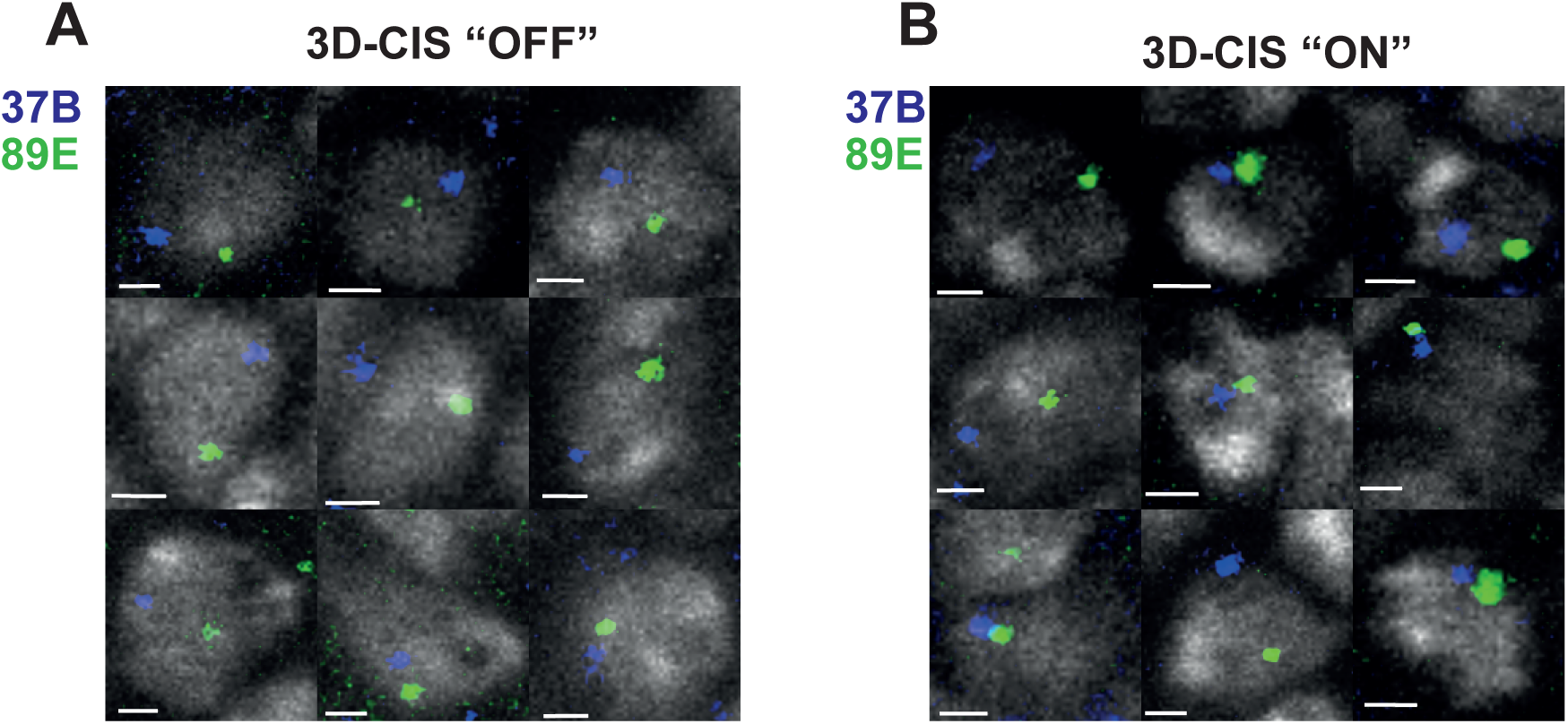
Activation of the 3D-CIS system induces chromatin contacts between Fab2L and *Fab-7*. **(A,B)** Micrograph galleries of randomly selected nuclei of the indicated genotypes in FISH-stained embryos. Nuclei are stained with DAPI in grey, the 37B locus surrounding the Fab2L transgene is stained in blue and the 89E locus surrounding the endogenous *Fab-7* is stained in green. Scale bar represents 1 μm. The quantification associated with these images is represented in Figure 5.

**Figure S9.**
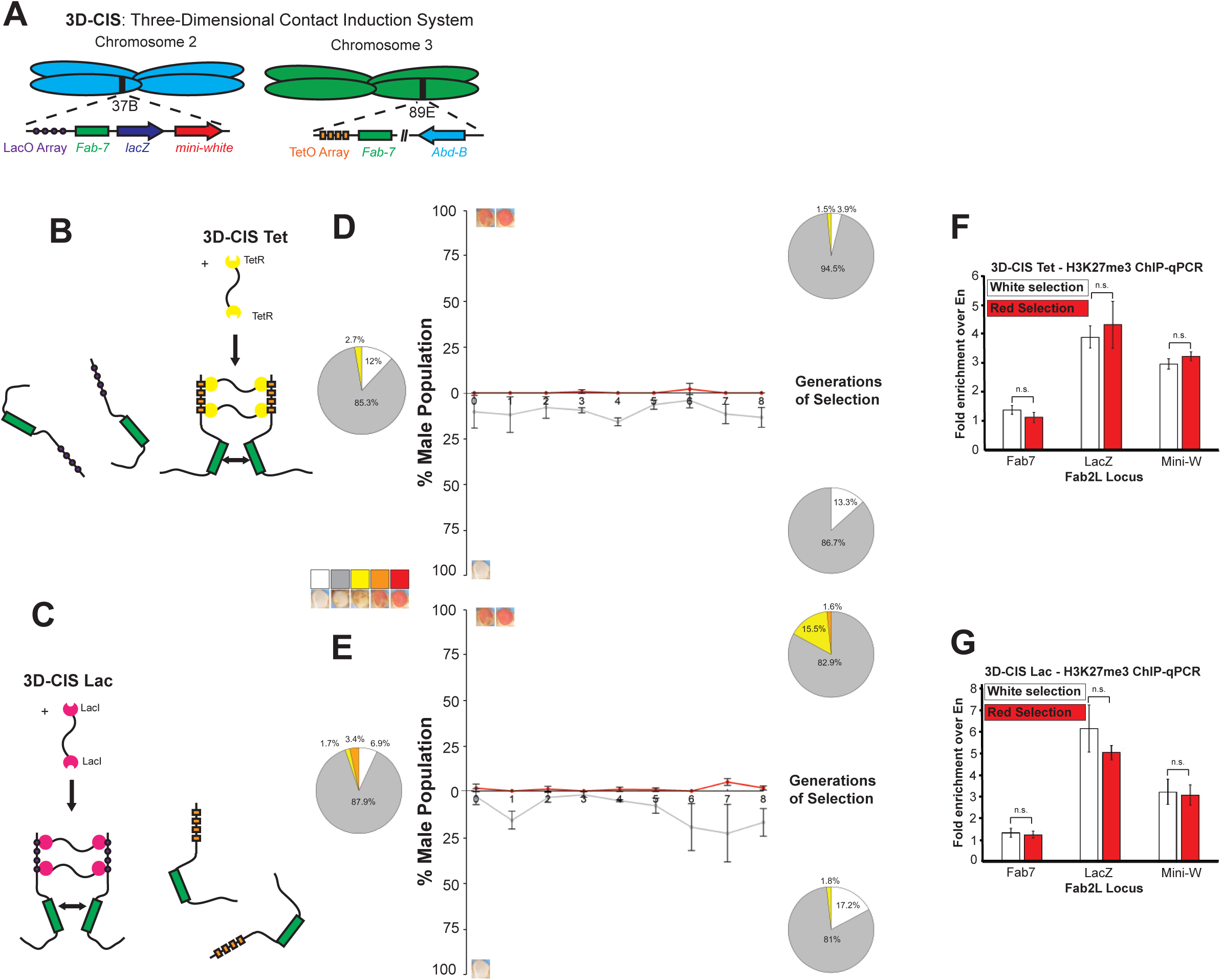
Promotion of chromatin contacts between homologous *Fab-7* alleles, rather than interchromosomal loci, using the 3D-CIS system does not lead to TEI establishment. **(A-C)** Schematic representation of the 3D-CIS Tet and 3D-CIS Lac systems, variations on the 3D-CIS, used as controls. Arrays of Lac and Tet operons are inserted next to the transgenic and endogenous *Fab-7* elements. Expression of a TetR-TetR or LacI-LacI fusion protein binding to only one array controls for the potential induction of contacts between homologous alleles rather than interchromosomal loci. **(D,E)** Results of selection for the most white or red-eyed flies in each generation of 3D-CIS Tet and 3D-CIS Lac flies in the “ON” state. Curves represent the percentage of males of Class 1 or Class 4+5 in the population across generations. Error bars are +/- SD of 3 independent repeats. Pie charts represent the phenotypic distribution of the eye colour within the population in the first and last generations of selection. **(F,G)** ChIP-qPCR assays agasint H3K27me3 performed in embryos of the indicated genotypes after selection towards white or red epiallele identity, at regions within the Fab2L transgene. Error bars represent +/- SEM of three independent repeats. Samples were normalised to engrailed as a positive control and compared to each other by the t-test (*p<0.05, **p<0.01, n.s. = not significant).

**Figure S10.**
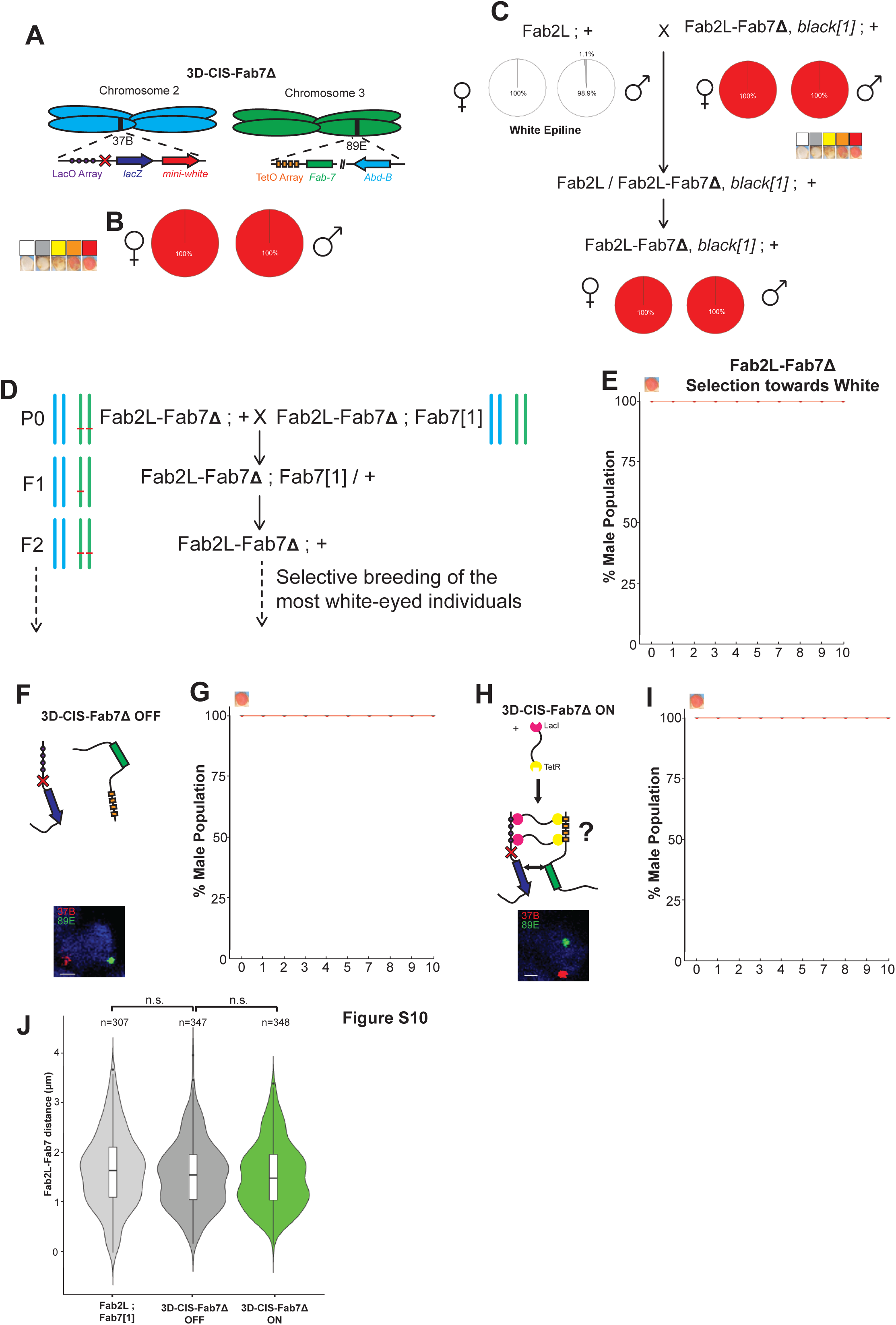
Artificial promotion of chromatin contacts is unable to significantly decrease inter-loci distance or trigger TEI in the absence of the *Fab-7* element. **(A,F,H)** Schematic representation of the 3D-CIS-Fab7Δ line in which the *Fab-7* element was deleted from the Fab2L transgene of the 3D-CIS line. **(B)** Phenotypic classification of eye colour of the 3D-CIS-Fab7Δ flies. **(C)** Paramutation crossing scheme and phenotypic distribution of the populations with the indicated genotypes and epiline identities. **(D)** Crossing scheme for the triggering of TEI in Fab2L-Fab7Δ, with diagrammatic representation of the copy number of the *Fab-7* element on chromosomes 2 and 3. **(E,G,I)** Results of selection for the most white-eyed flies in each generation after the cross or in the 3D-CIS system. Curves represent the percentage of Class 5 males in the population across generations. Error bars are +/- standard deviation (SD) of 3 independent repeats. **(J)** Violin plots representing the distance distributions of the 37B and 89E regions surrounding the two *Fab-7* elements as determined by FISH in the indicated genotypes. Distances were measured in stage 14-15 embryos in T1 and T2 segments. Distributions were compared using the t-test (n.s. = not significant, i.e. p>0.05).

**Figure S11.**
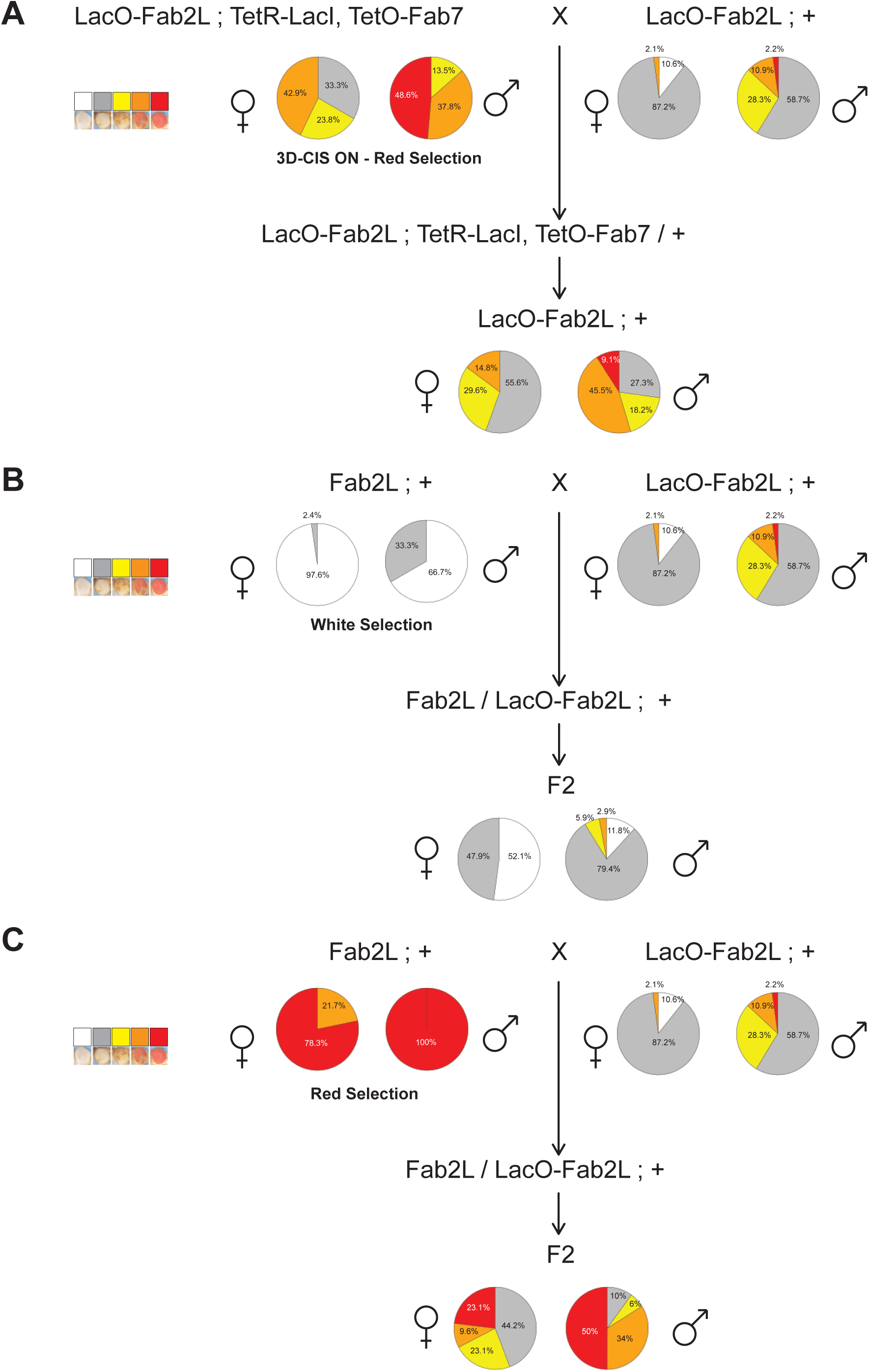
Altered epigenetic states remain stable in the absence of artificially-induced chromatin contacts. **(A-C)** Paramutation crossing schemes and phenotypic distribution of the populations with the indicated genotypes and epiline identities. Pie charts represent the phenotypic distribution of the eye colour within the population separated into five classes.

